# Seasonal temporal dynamics of marine protists communities in tidally mixed coastal waters

**DOI:** 10.1101/2021.09.15.460302

**Authors:** Mariarita Caracciolo, Fabienne Rigaut-Jalabert, Sarah Romac, Frédéric Mahé, Samuel Forsans, Jean-Philippe Gac, Laure Arsenieff, Maxime Manno, Samuel Chaffron, Thierry Cariou, Mark Hoebeke, Yann Bozec, Eric Goberville, Florence Le Gall, Loïc Guilloux, Anne-Claire Baudoux, Colomban de Vargas, Fabrice Not, Eric Thiébaut, Nicolas Henry, Nathalie Simon

## Abstract

Major seasonal community reorganizations and associated biomass variations are landmarks of plankton ecology. However, the processes determining marine species and community turnover rates have not been fully elucidated so far. Here, we analyse patterns of planktonic protist community succession in temperate latitudes, based on quantitative taxonomic data from both microscopy counts and ribosomal DNA metabarcoding from plankton samples collected biweekly over 8 years (2009-2016) at the SOMLIT-Astan station (Roscoff, Western English Channel). Considering the temporal structure of community dynamics (creating temporal correlation), we elucidated the recurrent seasonal pattern of the dominant species and OTUs (rDNA-derived taxa) that drive annual plankton successions. The use of morphological and molecular analyses in combination allowed us to assess absolute species abundance while improving taxonomic resolution, and revealed a greater diversity. Overall, our results underpinned a protist community characterised by a seasonal structure, which is supported by the dominant OTUs. We detected that some were partly benthic as a result of the intense tidal mixing typical of the French coasts in the English Channel. While the occurrence of these microorganisms is driven by the physical and biogeochemical conditions of the environment, internal community processes, such as the complex network of biotic interactions, also play a key role in shaping protist communities.

## 1. Introduction

Annual successions of species - and associated variations in biomass - are one of the classical hallmarks of plankton ecology in both marine and freshwater systems (Cushing, 1959; Margalef, 1978; Sommer et al., 1986; Winder & Cloern, 2010; Sommer et al., 2012). In temperate biomes, annual plankton biomass patterns classically involve some regularity in the form of a phytoplankton spring bloom (Sverdrup, 1953; Cushing, 1959; Margalef, 1978) that follows the increase of light availability in relation to a decrease in vertical mixing and nutrient availability, and provides food to grazers. The resulting spring peak of zooplankton leads to the decline of phytoplankton towards a mid-season biomass minimum while subsequent food limitation and fish predation controls zooplankton biomass. The sequence of planktonic taxa emerging along the course of this rhythmic phenomenon depends on regional geographical, ecological and biogeochemical specificities (e.g., coastal *versus* shelf *versus* oceanic conditions), but the overall annual reoccurrence of the same dominant species shows striking regularities in a given habitat. Such annual persistent patterns of species successions are well known for phytoplankton and zooplankton (Margalef, 1978; Modigh et al., 2001; Ribera d’Alcalà et al., 2004). In most regions, these seasonal cycles linked to plankton species phenologies have probably governed the evolution of life cycles and migratory behaviors of organisms ranging from the smallest fishes to whales and birds (Cushing, 1969; 1990; Longhurst, 1998). Identifying these temporal patterns and determining their principal environmental drivers are essential to reveal the mechanisms that drive species succession and that shape community composition, and to predict how climate change is likely to modify these patterns (Edwards & Richardson, 2003; Siano et al., 2021).

Predicting the consequences of environmental changes on the seasonal successions of plankton species is extremely challenging since not all mechanisms that produce the sequence of species along a seasonal cycle have been elucidated. Decades of research have emphasized the major role of physical factors (e.g., light and turbulence; Margalef, 1978; Towsend et al., 1992, 1994; Sommer et al., 2012; Barton et al., 2014) in pacing the annual oscillations of plankton biomass and diversity. These factors, contingent to the annual climate cycle and operating across various astronomic and geological time scales, would impose synchrony on the dynamics of phytoplankton biomass (Smetacek, 1985; Sommer et al., 1986; Cloern, 1996) in a similar way to terrestrial plants (Richardson et al., 2010; Craine et al. 2012). But seasonal successions are also an emergent property of the community dynamics, where the complex network of biotic interactions (e.g., predation, competition, parasitism, mutualism) and the species internal clocks - that are tightly set by the photoperiod - determine the self-organization and resilience of species assemblages for a specific time (Drake et al. 1990; Dakos et al. 2009; Logares et al., 2018). Self-organization processes selected by environmental filters, could be the major force shaping annual plankton successions. In other words, intrinsic community biological factors, including interactions within and between species and functional groups, could drive the stability of marine plankton through time. The idea that biodiversity buffers ecosystem changes against environmental variations (Tilman, 1999; Tilman et al., 2006; Loreau & Manzancourt, 2013) matches results obtained from manipulated microbiomes (Fernandez-Gonzalez et al., 2016) and theoretical studies (Dakos et al., 2009). It could explain the strong temporal relationship that links species richness and community-level properties (Cottingham et al., 2001; Loreau et al., 2001; Griffin et al., 2009).

Characterized by particularly high dispersal capabilities, large population sizes, and short generation time (Villarino et al., 2018), marine microorganisms represent about half of overall carbon biomass and play key roles in global biogeochemical fluxes (Falkowski et al. 2008; Bar-On and Milo, 2019). They are responsible for nearly all of the primary production and respiration occurring in the marine realm (Moran, 2015). While annual species successions have been hard to demonstrate for microorganisms (i.e., viruses, bacteria, archaea and protists; phototrophic and non-phototrophic microeukaryotes), especially for those with sizes under 10 µm which are difficult to identify under a microscope, the use of High Throughput Sequencing (HTS) and metagenomics approaches has shown that marine microbial communities exhibit clear annual patterns of species or Operational Taxonomic Units (OTU) successions (Fuhrman et al., 2006, 2015; Gilbert et al., 2012; Bunse & Pinhassi, 2017; Giner et al., 2018; Käse et al., 2020). Given the extraordinary roles of microscopic plankton in ocean ecology, being able to document their dynamics in space and time is of tremendous importance to predict future changes that will occur in the next decades.

Here, we report pluri-annual patterns of protists community dynamics at the Roscoff SOMLIT-Astan station, a coastal long-term time-series sampling site located off the French coast of the Western English Channel (WEC) (Fig. 1). The English Channel (EC) is an epicontinental sea which stands as a biogeographical crossroad between the warm-temperate Atlantic system and the cold-temperate North Sea and Baltic continental system of Northern Europe. Significant biological shifts, including species replacements or major changes in species abundances and distributions, have been documented in the English Channel since over a century in response to climate change and other anthropogenic drivers (Boalch et al., 1987; Southward et al., 2005; Mieszkowska et al., 2014). There are indications that current anthropogenic climate changes have already impacted pelagic and benthic compartments and affected the productivity of this shelf sea (see for example, Beaugrand et al., 2002; Genner et al., 2004; Hiscock et al., 2004; Southward et al., 2005, Widdicombe et al. 2010). The EC is a zone of high turbulence due to strong tidal currents. A seasonal thermocline, occurring from May to October, is only reported in its western entrance, offshore and along the UK coasts (Pingree and Griffiths, 1980). In these seasonally stratified waters sampled regularly by the Plymouth L4 Western Channel Observatory (Pingree and Griffiths, 1978), plankton temporal dynamics has been recently explored (Widdicombe et al., 2010; Edwards et al., 2013, Barton et al., 2020) and both seasonal and inter-annual changes in abundance were observed along with significant long-term changes in community composition and reorganization of plankton food web (Molinero et al., 2013; Reygondeau et al., 2015). The temporal dynamics of planktonic communities in the permanently well-mixed waters that characterize the French coasts of the WEC have been less intensively studied. In this region, the hydrodynamics is mostly driven by intense tidal currents and density gradients due to inflows of small rivers. These features are at the origin of strong physical and biogeochemical heterogeneity and of a mosaic of interconnected benthic and pelagic habitats (Dauvin et al., 2008; Delavenne et al., 2013; Gac et al., 2020). At the Roscoff time series station, the daily and seasonal biological cycles seem to maintain biogeochemical fluxes in steady state, for example in terms of CO_2_ fluxes (Gac et al., 2020), suggesting that the complex plankton community has the capacity to buffer environmental changes at the scale of at least a few years.

**Figure 1.**
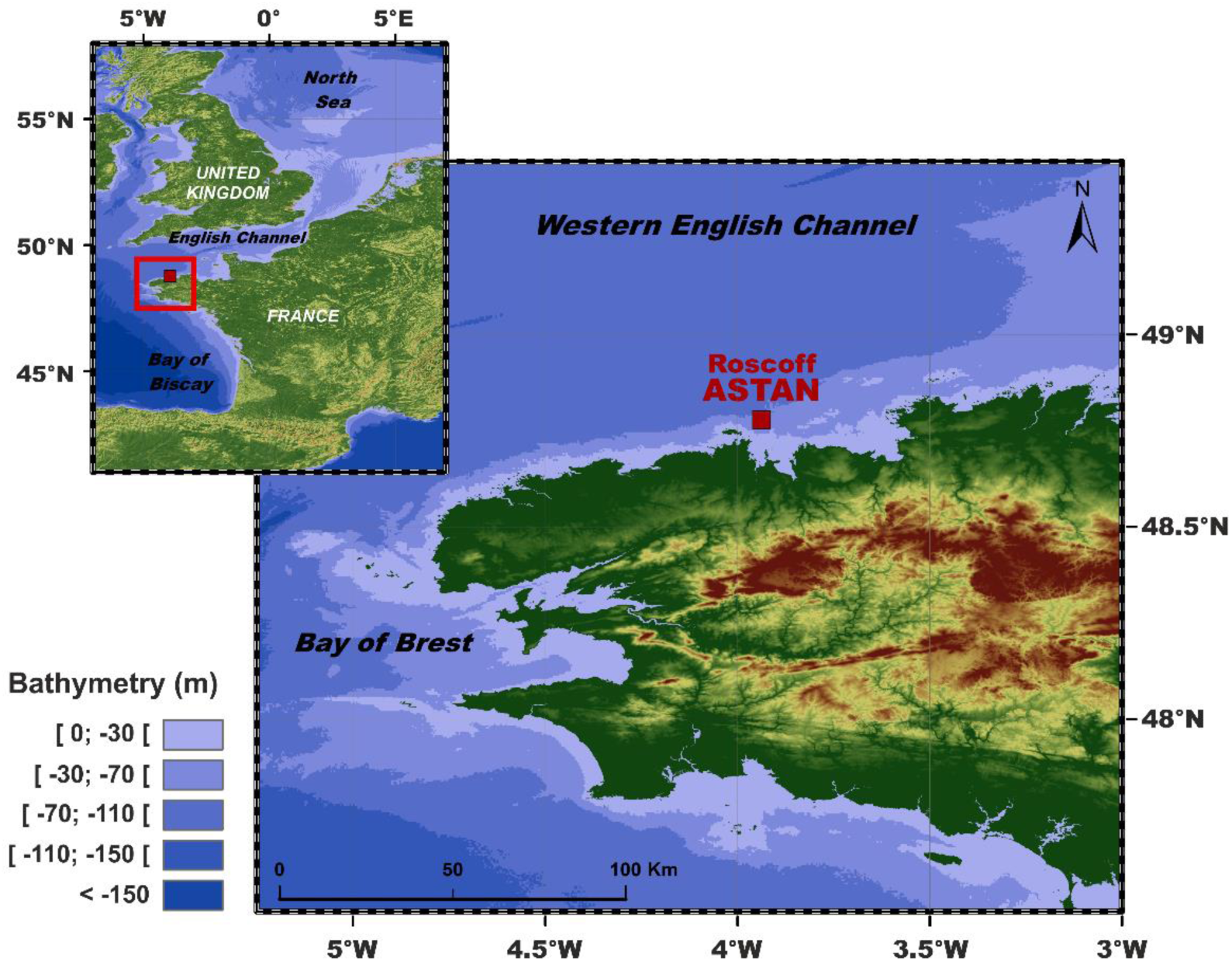
Location of the study area. The SOMLIT-Astan sampling station (48:46’49’’ N; 3:58’14’’ W) is located in the Western English Channel, 3.5 km from the coast. The water column at this site is 60 m deep and is never stratified due to intense tidal mixing. The site is strongly impacted by storms in Winter.

In this study, our aim was to (*i*) describe the seasonal dynamics of the protist eukaryotic community in this macro tidal coastal marine plankton ecosystem across multiple years, and (*ii*) explore how environmental factors interact with the dominant biotic compartment at different time scales. We analysed time-series of phytoplankton cell microscopy counts and DNA amplicons V4 sequence abundances generated from plankton samples collected over a period of 8 years (2009-2016). We unveil the protist community dynamics of this region in details, and provide new data on the dynamics of taxa that cannot be studied with classical microscopic techniques. Our results suggest that the temporal structure of environmental factors has an important influence on the seasonal dynamics of the community that exhibited recurrent successions over the 8-year period. Self-organization processes selected by environmental filters are suspected to be a major force shaping annual plankton successions.

## 2. Material and methods

### 2.1 Sampling station

The SOMLIT-Astan sampling station is located in the western English Channel, 3.5 km off Roscoff (Brittany, France) (60 m depth, 48°46’18” N–3°58’6” W, Fig. 1). In this coastal station, considered as representative of the French Western English Channel (Tréguer et al., 2014) intense tidal mixing (tidal range up to 10 m) and winds prevent summer stratification (Grall 1972, Martin-Jezequel 1983, Sournia et al. 1987). This station is also under limited influence by coastal freshwater inputs as nutrients loading from the adjacent rivers is rapidly diluted by tidal currents (L’Helguen et al. 1996, Wafar et al. 1983, Tréguer et al. 2014), but it is strongly impacted by the weather conditions that can be rough in the area with frequent gusts of wind and storms. Monitoring of the hydrology and phytoplankton at the SOMLIT-Astan station has been implemented in 2000 (Guilloux et al. 2013), and is currently operated in the frame of the SOMLIT (Service d’Observation en Milieu LITtoral, since 2000, http://somlit.epoc.u-bordeaux1.fr/) and PHYTOBS (PHYtoplankton OBServatory, since 2018) national monitoring programs. Plankton samples are collected bi-monthly at the SOMLIT-Astan station during high neap tide at surface (1 m depth) using a 5L Niskin bottle, as well as hydrological parameters measured during the same dates. For this study, the data corresponding to the period 2009 to 2016 were analysed.

### 2.2 Environmental data

Meteorological data (rainfall height, wind speed and direction, global radiation) and hydrological data (temperature, nutrient concentrations, chlorophyll-*a* biomass, particulate organic carbon and nitrogen, and suspended matter) for the period 2009 to 2016 were obtained from MétéoFrance (https://meteofrance.com/) and the SOMLIT program (https://www.somlit.fr/parametres-et-protocoles/), respectively. Mean daily tidal amplitude values were calculated from the water hourly heights available from the Service Hydrographique et Océanographique de la Marine (SHOM, https://data.shom.fr/), and used as a proxy of tidal mixing. Parameters related to light were obtained from the National Aeronautic and Space Administration (NASA, https://modis.gsfc.nasa.gov/data/dataprod/) and the National Oceanic and Atmospheric Administration (NOAA, https://coastwatch.pfeg.noaa.gov/). The average light received during the 8 days that preceded each sampling dates was calculated from PAR (photosynthetically available radiation, PAR8days, extracted from https://modis.gsfc.nasa.gov/data/dataprod/par.php). The diffuse attenuation coefficient for downwelling irradiance at 490 nm (Kd490), dependent on the availability of ratios of remote sensing reflectance (Rrs) in the blue-green spectral region (e.g., 490 - 565 nm) was also extracted (from https://modis.gsfc.nasa.gov/data/dataprod/kd_490.php). The North Atlantic Oscillation index (NAO, Hurrell, 1995; Trigo et al., 2002) that influences the local meteorological conditions, was obtained from NOAA (https://www.ncdc.noaa.gov/teleconnections/nao/). Protocols used for the hydrological parameters by the SOMLIT are summarized below (Gac et al., 2020, 2021). Seawater temperature (T°C) was measured *in situ* using a Sea-bird SBE19+ CTD profiler with an initial accuracy of +/- 0.005°C. Discrete salinity samples were measured on a portasal salinometer with a precision of 0.002. Nutrient concentrations (NO_3_^-^, NO_2_^-^, PO_4_ and SiO_4_^2-^) were determined using an AA3 auto-analyser (Seal Analytical) following the method of Aminot and Kérouel (2007) with an accuracy of 0.02 µmol L^-1^, 1 nmol L^-1^, 1 nmol L^-1^ and 0.01 µmol L^-1^ for NO_3_^-^, NO_2_^-^, PO_4_ and SiO_4_ , respectively. Ammonium (NH_4_ ) concentrations were determined using the indophenol blue method of Koroleff (1969). To determine chlorophyll-*a* concentrations (Chl*-a*), 0.5 L of seawater were filtered onto glass-fibre filters (Whatman GF/F) and immediately frozen. Samples were extracted in 5 mL of acetone, acidified with HCl and Chl*-a* concentrations, and were measured using a fluorometer (model 10 analog fluorometer Turner Designs), according to EPA (1997), with an estimated accuracy of 0.05 µg L^-1^. Protocols used to measure the biomass of particulate organic carbon (POC), particulate organic nitrate (PON) and suspended matter (MES) are described in the SOMLIT website.

### 2.3 Phytoplankton microscopic counts

Samples (250 mL) of natural seawater intended for the acquisition of microscopic counts were preserved with acid Lugol’s iodine (Sournia, 1978, Guilloux et al. 2013), stored in the dark, and further processed between 15 days and up to 1 year after sampling. Lugol’s iodine was added either back in the lab 1.5 to 2h after sampling or onboard immediately after sampling. Cell counts were obtained from sub-samples that were gently poured into 50 mL composite settling chamber (HYDRO-BIOS, Kiel), according to the standard Utermöhl settlement method (Sournia, 1978; Guilloux et al., 2013). For some winter samples with low cell densities, 100 mL settlement chambers were used. Counts and identification of taxa were performed under an inverted light microscope (Leica DMI 300) at 200x and 400x magnification. References used for species identification included Tomas (1997), Throndsen et al. (2007), Hartley et al. (1996), Kraberg et al. (2010), Hoppenrath et al. (2009), Horner (2002) and the Plankton*Net Data Provider (http://www.planktonnet.eu/). Taxonomic assignation was achieved to the highest taxonomic rank that we could be reached, at species level when possible. Raw microscopic counts were regularly stored in a local MS-Access database and uploaded in the RESOMAR PELAGOS (http://abims.sb-roscoff.fr/pelagos/) national database. The morphological taxa contingency table was carefully examined to detect inconsistencies (e.g., abrupt changes in cell counts over the time series), and taxa for which identification was uncertain were grouped into broader taxonomic categories. For example, *Fragilaria* and *Brockmaniella* or *Cylindrotheca closterium* and *Nitzschia longissima* which are difficult to distinguished between each other, were considered in association in the same group of microscopic counts. The final morphological dataset (DOI: 10.5281/zenodo.5033180) consisted of counts of 146 taxonomical entities (taxa larger than 10µm in size) across 185 dates from 2009 to 2016.

### 2.4 Protists DNA metabarcodes

For the generation of DNA metabarcoding data, natural seawater from the Niskin bottle was transported to the laboratory in a 10L Nalgene bottle and a volume of 5L was collected onto 3µm polycarbonate membranes (47mm, Whatman; with the exception of May 25^th^ 2010, when the sample was collected onto a 0.22µm sterivex filter, PVDF, Millipore). Filters were preserved in 1.5mL of lysis buffer (Sucrose 256g/L, Tris 50mM pH8, EDTA 40mM) and stored at -80°C until further processing. A total of 185 samples were collected between 2009 and 2016.

#### 2.4.1 Plankton DNA extraction, PCR amplification, and metabarcode sequencing

The DNA extraction and generation of metabarcodes were performed using the exact same procedure for all samples. Samples were first incubated 45min at 37°C with 100µL lysozyme (20mg/mL), and 1h at 56°C with 20µL proteinase K (20mg/mL) and 100 µL SDS 20%. Nucleic acids were then extracted using a phenol-chloroform method (Sambrook et al. 1989), and purified using silica membranes from the NucleoSpin® PlantII kit (Macherey-Nagel, Hoerdt, France). DNA was eluted with 100µL Tris-EDTA 1x pH8 buffer and quantified using a Nanodrop ND-1000 spectrophotometer and a Qubit 2.0 Fluorometer instrument with dsDNA HS (High Sensitivity) assay (ThermoFisher Scientific, Waltham, MA). Total DNA extracts were then used as templates for PCR amplification of the V4 region of the 18S rDNA (∼380 bp) using the primers TAReuk454FWD1 (5’-CCAGCASCYGCGGTAATTCC-3’, *S. cerevisiae* position 565-584) and TAReukREV3 (5’-ACTTTCGTTCTTGATYRA-3’, *S. cerevisiae* position 964-981) (Stoeck et al. 2010) that target most eukaryotic groups. The forward primer was linked to a tag, and both primers were adapted for Illumina sequencing. PCR reactions (25 μl) contained 1x Master Mix Phusion High-Fidelity DNAPolymerase (Finnzymes), 0.35 μM of each primer, 3% dimethylsulphoxide and 5 ng of DNA. Each DNA sample was amplified in triplicates. The PCR program had an initial denaturation step at 98°C during 30 s, 10 cycles of denaturation at 98°C, annealing at 53°C for 30 s and elongation at 72°C for 30 s, then 15 similar cycles but with 48°C annealing temperature, and a final step at 72°C for 10 min. Polymerase chain reaction triplicates were pooled, and purified and eluted (30 μl) with NucleoSpin Gel and PCR Clean-Up kit (Macherey-Nagel, ref: 740770.50 and 740770.250), and quantified with the Quant-It PicoGreen double stranded DNA Assay kit (ThermoFisher). About 1 μg of pooled amplicons were sent to Fasteris (www.fasteris.com, Plan-les-Ouates, Switzerland) for high throughput sequencing on a 2×250bp MiSeq Illumina. Sequences were obtained in five separate runs. Overall, ∼ 7million unique sequences were obtained for a total of 185 samples collected over the 8 years (> 3µm).

#### 2.4.2 Reads quality filtering and clustering

Generation of 18S V4 rDNA Operational Taxonomic Units (OTUs) from the raw sequencing reads and their assembly into a contingency table was obtained according to the following pipeline (https://nicolashenry50.gitlab.io/swarm-pipeline-astan-18sv4). The paired-end fastq files were demultiplexed and PCR primers were trimmed using Cutadapt v2.8 (Martin, 2011). Reads shorter than 100 nucleotides or untrimmed were filtered out. Trimmed paired-end reads were merged using the fastq mergepairs command from VSEARCH v2.9.1 (Rognes et al., 2016) with a minimum overlap of 10 base pairs. Merged reads longer than 200 nucleotides were retained and clustered into OTUs using Swarm v2.2.2 with *d* = 1 and the *fastidious* option (Mahé et al., 2014, 2015). The most abundant sequence of each OTU is defined as the representative sequence. OTUs with a representative sequence considered to be chimeric by the uchime_denovo command from VSEARCH or with a quality per base below 0.0002 were filtered out. Samples with a low number of reads were re-sequenced, for these cases, only the readset with the highest number of reads was kept. Finally, OTUs which appeared in less than 2 samples or with less than 3 reads were discarded (de Vargas et al., 2015).

#### 2.4.3 Taxonomic assignations

The V4 region was extracted from the 18S rDNA reference sequences from PR2 v4.12 (Guillou et al., 2013) with Cutadapt, using the same primer pair as for the PCR amplification (maximum error rate of 0.2 and minimum overlap of 2/3 the length of the primer). The representative sequences of each OTU were compared to these V4 reference sequences by pairwise global alignment (usearch_global VSEARCH’s command). Each OTU inherits the taxonomy of the best hit or the last common ancestor in case of ties. OTUs with a score below 80% similarity were considered as unassigned (Mahé et al., 2017; Stoeck et al., 2010). In this study, focusing on the ecology of protists, only OTUs assigned to protist lineages (eukaryotes which are not Metazoa, Rhodophyta, Phaeophyceae, Ulvophyceae or Streptophyta) were considered. The final dataset (filtered OTU table, available at DOI:10.5281/zenodo.5032451) contained 185 samples with a total of ∼12.7 million sequence reads and 15,271 OTUs affiliated to protist taxa. Assignation of the dominant OTUs (e.g., based on abundance and occurrence) was checked and refined manually by BLASTing them (https://blast.ncbi.nlm.nih.gov/Blast.cgi) against the SILVA reference database (https://www.arb-silva.de/). The origin and assignations of the best blast sequences (most of which were 100% similar to our sequences) and of the corresponding strains or isolates were carefully examined before taking the final taxonomic assignation decision (Table S1).

### 2.5 Statistical analyses

A summary of all analyses performed for the metabarcoding dataset is illustrated in Fig. S1 and detailed procedures are available in GitLab (https://gitlab.com/MariaritaCaracciolo/roscoff-astan-time-series). The analyses performed on the morphological dataset were similar except that absolute abundance (cell counts) were used for all analyses (no rarefaction step prior to the calculation of species richness, Shannon Diversity Index and Jaccard similarity).

#### 2.5.1 Alpha and Beta diversity

Standard alpha diversity metrics (Shannon Diversity Index and species richness) and beta diversity metrics (Jaccard similarity index and Bray-Curtis similarity index; Krebs, 1999; Legendre & Legendre, 1998) were calculated for both the morphological and metabarcoding datasets in order to analyse temporal changes in the composition and structure of the protist communities Random subsampling (rarefaction) was used for the metabarcoding dataset prior to the calculation of alpha diversity metrics and for the calculation of the Jaccard similarity index in order to account for differences in sequencing depth (i.e. total number of reads generated for a sample). Hellinger transformed data (Legendre & Gallagher, 2001) were used for the calculation of Bray-Curtis dissimilarities (d = 1 – S, where d is dissimilarity and S is similarity between samples). The transformation is necessary for metabarcoding data where only relative abundance is meaningful.

### 2.5. Detecting the temporal structure of plankton protist community

In order to detect the temporal structure of the communities, we used distance-based Moran’s eigenvector maps (dbMEM) (Legendre & Gauthier, 2014). This method has the potential to detect temporal structures produced by the species assemblage itself (through auto-assemblage processes or autogenetic succession that involve species interactions, Connell and Slatyer, 1977; Reynolds, 1984; McCook et al., 1994) provided that all influential variables have been included in the analysis (Legendre and Gauthier, 2014). The dbMEM eigenfunctions were computed from a distance matrix of the time separating observations, truncated at a threshold corresponding to the largest time interval (lag= 44 days) (Legendre & Gauthier, 2014). A forward selection procedure implemented in the package adespatial (“forward.sel” function; Dray et al., 2018) was used to identify significant dbMEM. Among the generated positive and negative dbMEM eigenfunctions (n=55 and n=129, respectively), only 52 positive dbMEM were retained for the metabarcoding dataset and 47 for the morphological dataset and used as explanatory variables for a redundancy analysis (RDA; Ter Braak, 1994). This analysis consists in a series of regressions performed on community matrices, i.e., OTU read abundance (n=15,271) or species cell counts (n=146) data. Only OTUs present in at least 10 out of the 185 total samples were retained and the data were Hellinger-transformed in order to (*i*) avoid overweighting rare species and (*ii*) be able to use Euclidean distances that allow to compute RDA (Legendre & Gallagher, 2001). Significant linear trends were then removed by computing the residuals, and Anova-like tests (with 999 permutations; Legendre, Oksanen & Braak, 2013) were implemented on the RDA to assess the significance of each constrained axis (p value < 0.05). To calculate the proportion of the variance explained by the significant axes, the adjusted R^2^ of the RDA result was used. Variance partitioning analyses allowed to filter out the variations due to temporal structures, or autocorrelation, which accommodate the use of statistical tests to further assess which environmental variables can influence community dynamics and species composition. All parameters were first tested for collinearity, then successively used in a forward selection to identify those significant to be tested for the study. To interpret temporal variations, we calculated Spearman’s rank correlation coefficients between the environmental parameters and the eigenvalues of the first three axes of the RDA.

All statistical analyses were performed using the R environment (R version 4.1.0, R Development Core Team, 2011). The R package “vegan” (Oksanen et al., 2013) and “data.table” (Dowle and Srinivasan, 2018) were used to analyse frequency count data, diversity, and to compute variance partitioning. The dbMEM analyses were performed using the packages “ade4” (Dray and Dufour, 2007), “adespatial” (Dray et al., 2018), “ape 5.0” (Paradis & Schliep, 2019) and “spdep” (Bivand & Wong, 2018). All figures were made with “ggplot2” (Wickham, 2009).

## 3. Results

### 3.1 Seasonal ecosystem dynamics at the SOMLIT-Astan station

At the SOMLIT-Astan time-series station (Fig. 1), as expected in temperate marine waters, both the hydrological parameters and phytoplankton biomass displayed clear seasonal patterns over the 8-year period (2009-2016, Fig. 2). In this tidally mixed environment, mean monthly temperatures varied from 9.8 (in March) to 15.7°C (in August). Mean monthly salinity ranged between 35.1 and 35.4 (from spring to autumn). Seasonal changes in chlorophyll *a* (Chl-*a)* concentration were characterized by broad summer maxima (from June to August) and large inter-annual variability (Fig. S2). From 2009 to 2016, mean monthly values were recorded between 0.4 and 1.5 µg L^-1^ (in December and July, respectively), and seasonal variations were synchronous with PAR (5.3 to 48.1 E m^-2^ day^-1^). Mean monthly minima in the main macronutrient concentration (PO_4_^3^, SiO_4_^2-^ and NO_2_^-^) that sustain phytoplankton production were recorded in summer, when phytoplankton biomass was high; however, macronutrients were never completely depleted (Fig. 2). Annual oscillations of pH were also recorded with minima in autumn. Although sampling occurred consistently during high neap tides, a clear biannual rhythm was detected in the mean monthly tidal amplitudes, which varied between 3.1 and 4.2 m with the highest mean values in late spring (May) according to the yearly change in the obliquity of the Earth’s Equator. From 2009 to 2016, all parameters exhibited large inter-annual variations and no significant decadal trend was detected (Fig. S2).

**Figure 2.**
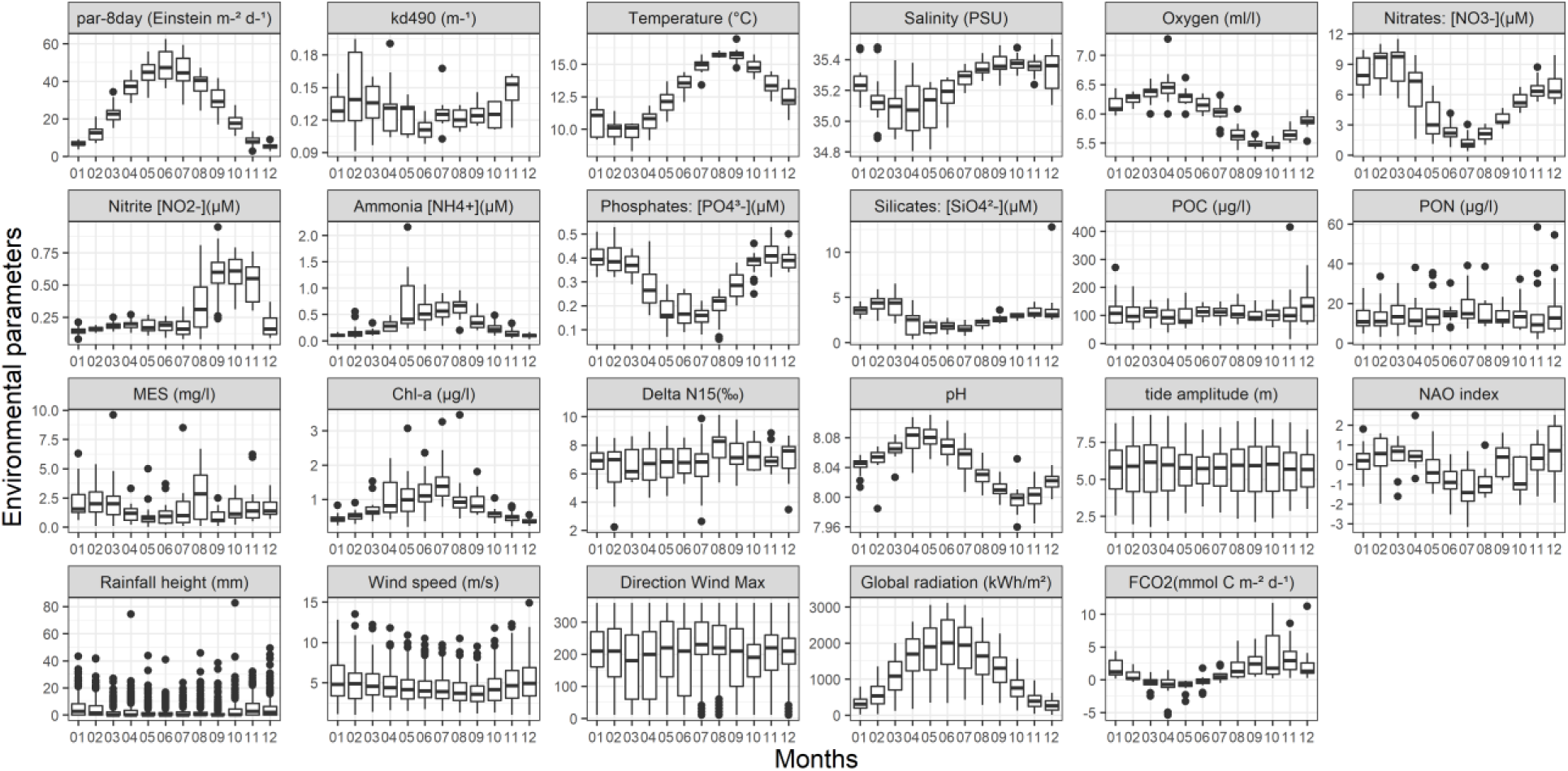
Monthly variations of the hydrological and meteorological parameters at the SOMLIT-Astan time-series station in the period 2009-2016. Sampling was carried out at high neap tides. PAR is the photosynthetically available radiation calculated as the average light received during the 8 days before sampling. Kd490 is intended as the diffuse attenuation coefficient for downwelling irradiance at 490 nm. Interannual variations of all parameters presented in this figure can be found in figure S2.

The protists community structure also showed clear seasonal patterns according to changes in alpha and beta diversity calculated from our morphological (only phytoplankton > 10 µm) and metabarcoding (all protist 18S V4rDNA OTUs > 3 µm) datasets (Fig. 3): minimal Shannon diversity was recorded in spring and summer, when Chl-*a* biomass was the highest, and maximum values in winter (Fig. 3A, B). This seasonal pattern, which is related to changes in the seasonal pattern of species dominance, was consistent among taxonomic groups although variations were encountered in the exact timing of the monthly minima of some of the phyla or classes distinguished using metabarcoding (Fig. S3). For groups such as the Cercozoa, an opposite signal was recorded (Fig. S3), with relatively high (low) Shannon’s diversity in spring and summer (winter). Taxa such as the MOCH-4 (marine Ochrophytes without cultured representatives), Perkinsea or Raphidophyceae were recorded almost exclusively during winter (Fig. S3).

**Figure 3.**
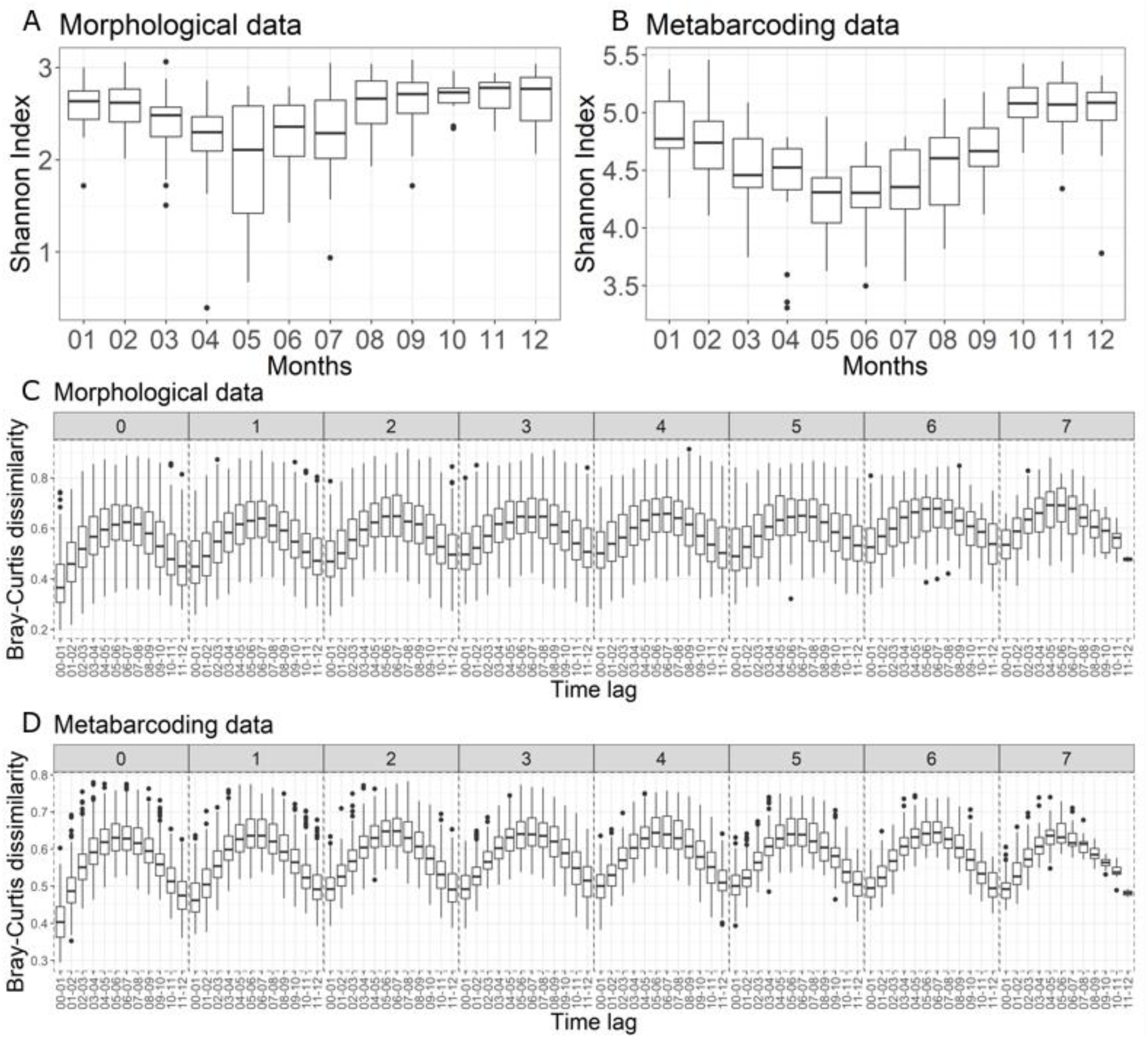
Changes in alpha and beta diversity calculated for the protist assemblages over the period 2009-2016 at the SOMLIT-Astan sampling station. (**A**, **B**): Seasonal variations in the Shannon Indexes calculated for the period 2009-2016. (**C**, **D**): Interannual recurrence of protist communities shown by the variations in the Bray-Curtis dissimilarity index between samples collected along the 2009-2016 period, as a function of increasing lag between sampling dates. The lag values between samples, for each box plot correspond to a number of years (facet labels, from 0 to 7) plus a number of months (x axis of each facet, expressed as ranges). For example, the lag between samples considered for the first box plot is 0 years and 0 to 1 months and the lag between samples considered for the last box-plot in 7 years and 11 to 12 months. Panels (**A**) and (**C**) are based on the morphological dataset (cell counts) while graphs (**B**) and (**D**) are based on the metabarcoding dataset.

The variations in the Jaccard and Bray-Curtis dissimilarities - calculated based upon the morphological and the metabarcoding datasets along temporal distances between samples - not only confirmed the strong seasonality in the structure of the community, but also suggested gradual replacements of taxa along the year and recurrence in the annual sequence of taxa over 8 years (Fig. 3B, D). The rates of changes in these similarities also showed clear temporal variations for both datasets and appeared to follow a biannual rhythm, with relative minima (maxima) in February-March and October (in May-July and December-January) (Fig. S4). A higher variability was recorded for the morphological dataset, with a decrease in similarity over time.

### 3.2 Overall composition of the protists assemblages in coastal mixed environments

Based on microscopy counts of plankton > 10 µm, diatoms were clearly the dominating group all year round and over the study period (86.5% and 74.4% of all cell counts and taxonomic entities distinguished, respectively Fig. 4A, C). Dinoflagellates covered another 7.1 % of all cells enumerated and accounted for 15.7% of total taxa richness. Ciliates and haptophytes (more precisely Oligotrichea and Prymnesiophyta) accounted for 2.4 and 2.1 % of all cell counts. The other groups such as Undetermined_sp., Raphidophyceae, Dictyochophyceae, Euglenophyceae, Pyramimonadophyceae, Xanthophyceae, Prasinophyceae, Undetermined_Chlorophyta, accounted each for 1% or less than 1% (Fig. 4A). Each of these groups accounted for < 3% of the total number of morphological entities (Fig. 4C).

**Figure 4.**
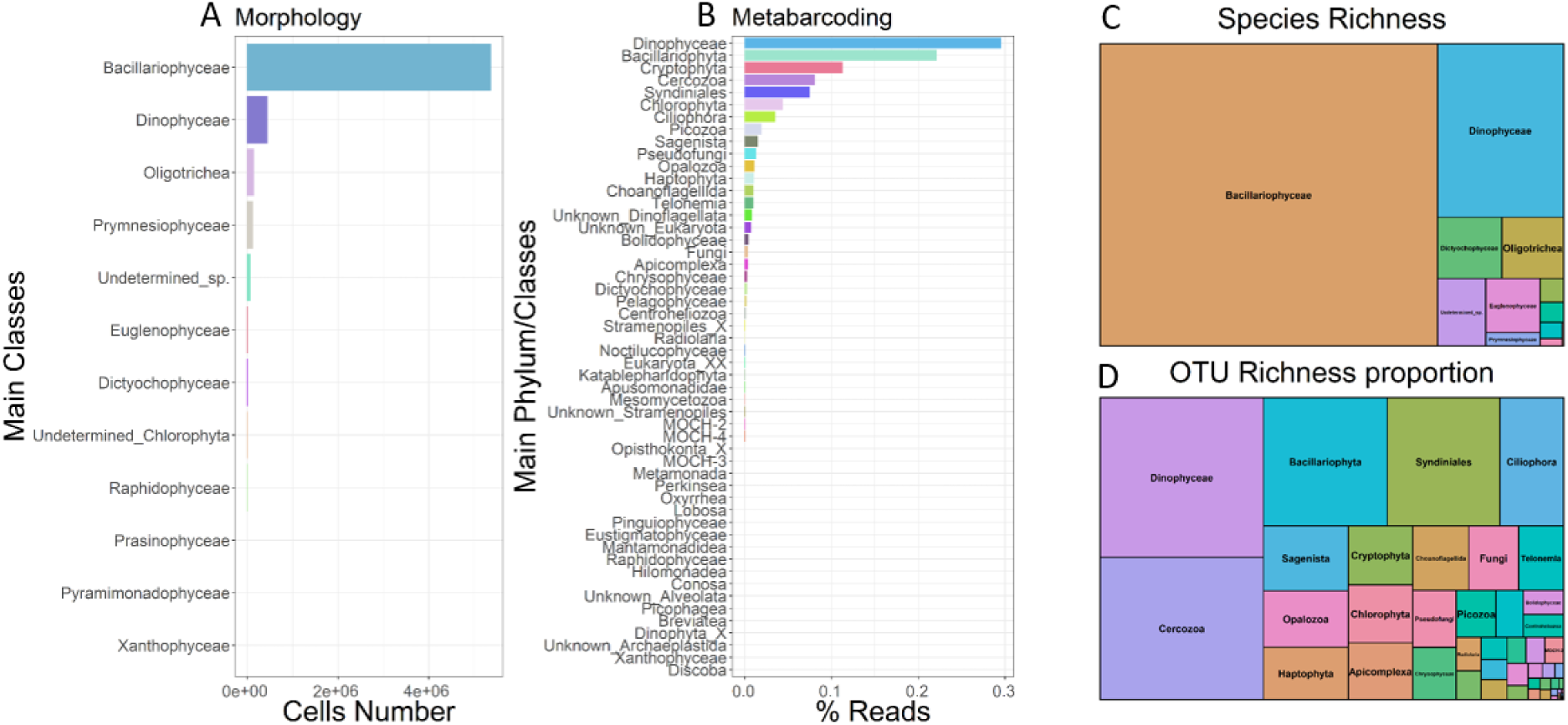
Low-taxonomic resolution contribution of protists at the SOMLIT-Astan time-series station over the period 2009-2016. Abundance of (**A**) the 12 main phytoplankton classes for the morphological dataset; and (**B**) the 52 main phyla - or classes – calculated from the metabarcoding dataset. The tree maps show the overall contributions of the main phyla or classes to the (**C**) total species or (**D**) OTU richnesses.

Our metabarcoding approaches based on total DNA extracts from plankton >3µm uncovered a much wider diversity spectrum. A first-order, low-resolution taxonomic assignation of all OTUs revealed the prevalence of Dinophyceae and diatoms in terms of reads numbers over the whole study period (29.6 and 22.1% of all reads, Fig. 4B, S5). Cryptophyta, Chlorophyta and Haptophyta that are primarily photosynthetic phyla accounted for 11.3%, 4.4% and 1.1% of all reads counts, while the heterotrophic Cercozoa, Syndiniales and Ciliophora, made up 8.1%, 7.5% and 3.5% of all read counts, respectively (Fig. 4B). The contributions of the other eukaryotes, including Picozoa, Sagenista, Pseudofungi, Opalozoa, Choanoflagellida, and Telonemia, were lower (< 2% of total reads, Fig. 4B). In terms of OTU richness, the picture was slightly different since Dinophyceae and cercozoans appeared as the first and second most diverse groups (18.6 and 16.6% of all OTUs, Fig. 4D), followed by diatoms, Syndiniales and Ciliophora (11.4, 10.3 and 5.8% of all OTUs). OTU richness from Sagenista (bicoecea and labyrinthulids), Opalozoa, Haptophyta, Cryptophyta, Chlorophyta, Apicomplexa, Choanoflagellida, Fungi and Telonemia ranged from 3.9% (Sagenista) to 2.1% (Telonemia) of the total number of OTUs. Other less diverse taxa belonging to 53 classes (e.g., Pseudofungi, Chrysophyceae, Picozoa, Dictyochophyceae, Bolidophyceae, Centroheliozoa, Radiolaria; see Fig. 4B for the complete list of the 52 classes) accounted for less than 2% of all OTUs (Fig. 4D).

Clear seasonal variations were encountered at phylum or class levels for absolute cell abundances (Fig. S5A). The abundances of diatom cells (>10um) generally peaked in late spring and summer, while dinoflagellates maximal abundances were observed in late summer. Important inter-annual variations were recorded in both the timing and intensity of the annual peaks, however. For the Prymnesiophyceae, the interannual variations were especially high, with exceptional developments of Haptophytes (corresponding to *Phaeocystis globosa* blooms) in spring 2012. Seasonal and inter-annual variations were also observed when contributions to total DNA reads abundances were examined, with maximal contributions of diatoms and Dinophyceae in spring and summer, respectively, and of Cryptophyta and Chlorophyta in summer and autumn, respectively. The contribution of Cercozoa and Syndiniales (and other primarily heterotrophic, parasitic or saprotrophic groups such as the Ciliophora, Picozoa, Opalozoa and Sagenista) started to increase in early winter and were high during the first months of the year (Fig. S5B).

### 3.3 Annual successions of protists at high taxonomic resolution

Given the clear annual recurrence of morphological and molecular taxa detected with beta diversity analyses (Fig. 3), we dug into seasonal variations of the protist assemblages at a finer taxonomic scale. First, to integrate the broadest taxonomic diversity including the smallest taxa, we used the metabarcoding dataset to calculate mean monthly relative abundances of all OTUs and select the 10 most abundant taxa for each month. This resulting list of 32 OTUs, due to the same OTUs being dominant in several months, contributed to 51.5% of all reads over the study period (Fig. 5A), and included diatoms, Dinophyceae, Cryptophyta, Cercozoa, Syndiniales, as well as a Chlorophyta, a Picozoa, a MAST and a Fungi. Sequences of both photosynthetic armored (*Heterocapsa*) and heterotrophic naked (e.g. *Warnowia* and *Gyrodinium*) dinoflagellates dominated the sequences pools all year round. The nanoplanktonic Cryptophytes *Teleaulax amphioxeia* (*= Plagioselmis prolonga*)*, T. gracilis* and *T. acuta* (all described as photosynthetic) and the green picoplanktonic algae *Ostreococcus lucimarinus* also appeared as dominant taxa. The sequences of several parasitic taxa such as the cercozoan *Cryothecomonas*, the dinoflagellates *Haplozoon* and Syndiniales, and the fungi *Parengyodontium* also showed high prevalence (Fig. 5A). Diatom OTUs identified as dominating the protist communities were assigned to *Minidiscus comicus, Minidiscus variabilis* and *Guinardia delicatula,* and to the genera *Thalassiosira* and *Arcocellulus* or *Minutocellus.* Although rather consistent over the 8 years, the temporal sequence showed important inter-annual variations (Fig. S6): for example, the relative contribution of reads assigned to the parasitic *Cryothecomonas* sp. and *C. linearis* were particularly prominent during the winters 2012 and 2013, and in July 2013 and 2015, respectively (Fig.S6A). Reads assigned to *Picozoa judraskeda* appeared only in 2016.

**Figure 5.**
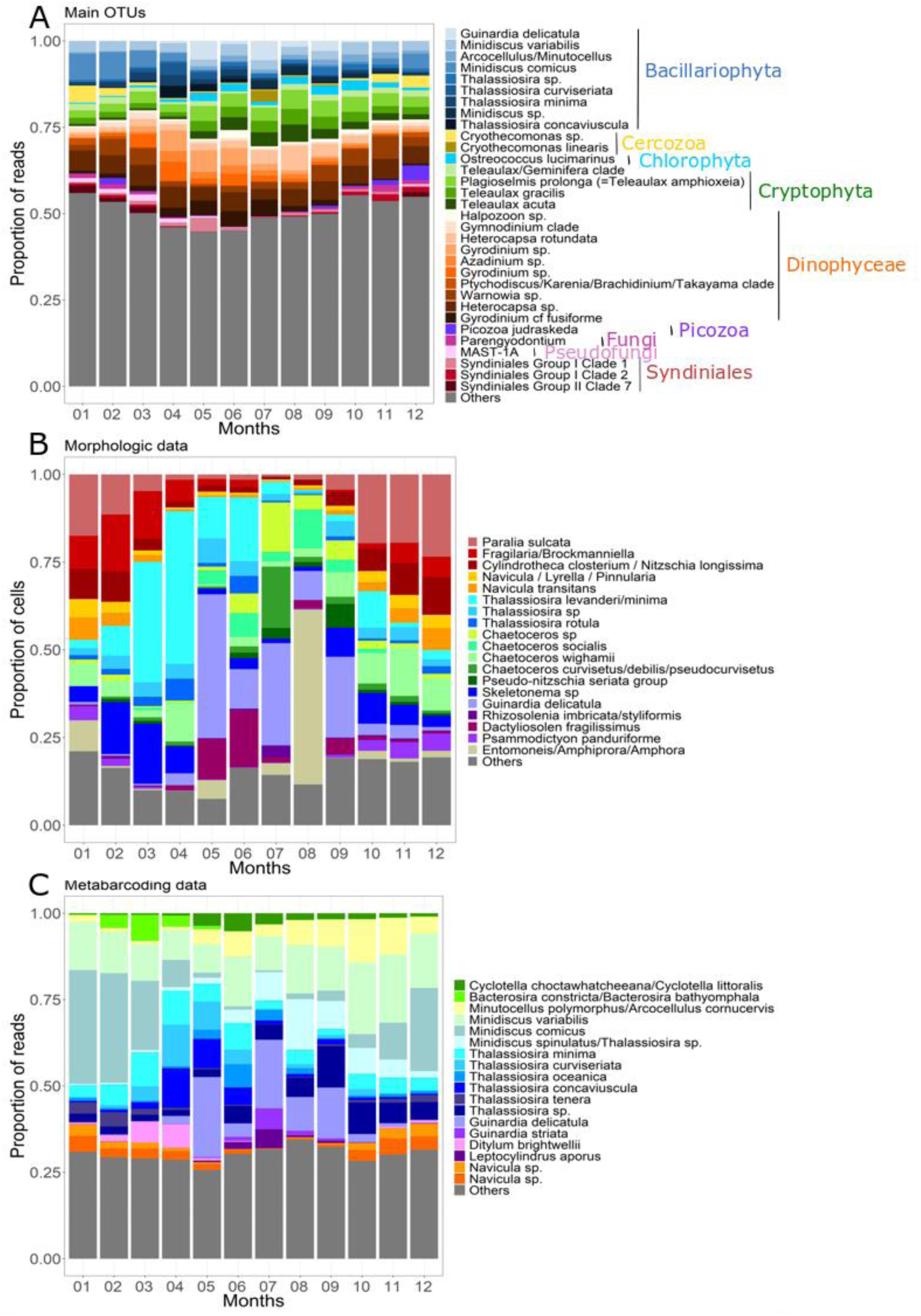
Typical seasonal variations of the dominant OTUs and overall contribution of the major diatoms species to the protist assemblage at the SOMLIT-Astan sampling station over the period 2009-2016. The histograms show the contributions (**A**) to total DNA reads abundance of the 32 dominating OTUs (accounting for 51.5% of all reads), (**B**) of the main diatoms to total diatoms abundances (microscopy count of plankton >10um) and (**C**) of the main diatoms to total diatom reads abundances. All microscopy counts and OTUs were assigned at the highest taxonomic level. Species selected were the 10 most abundant (5 for diatoms) for at least one month, taking into account mean monthly abundances.

A closer examination of the diatoms species dynamics was achieved by calculating mean monthly relative abundances of diatoms OTUs or cells and selecting the 5 most abundant taxa for each month in the microscopic and metabarcoding datasets (Fig. 5B, C). The resulting pools of cells or reads selected accounted for >75% and 70% of all counts/reads in these two datasets, respectively. In microscopic counts, autumn and winter assemblages were clearly dominated by species or genera with benthic affinities such as *Paralia* sp. and the pennate morphological combinations corresponding to *Fragilaria*/*Brockmaniella* and *Cylindrotheca closterium* /*Nitzschia longissima* (Fig. 5B). These taxa were replaced, from mid-winter to early spring, by colonial genera with pelagic affinities and in particular by *Thalassiosira* spp. (with *Thalassiosira levanderi/minima* reaching mean abundances of ∼534 cells.L^-1^ [35.83% of counts] in April) and *Skeletonema* spp. followed by *Dactyliosolen fragilissimus* all along spring. The dominant species in late spring and summer was *Guinardia delicatula* with the highest mean monthly abundances recorded in May and July (with ∼530 cells.L^-1^ for both months, 43% and 26.16% of diatoms counts, respectively). The contribution of the genus *Chaetoceros* was significant from spring until early winter (with *C. curvisetus/debilis/pseudocurvisetus and C. wighamii* showing relative high contributions in July and in winter, respectively). This picture of the mean yearly sequence of diatoms appeared rather resilient over the period 2009-2016, but inter-annual variations were apparent, with exceptional blooms of *Skeletonema* in early spring in 2011, 2013 and 2014, and *Chaetoceros socialis* in July 2014. The contribution of the benthic diatoms associated to the genera *Entomoneis/Amphiprora/Amphora* was exceptionally high in 2011.

The analysis of the genetic dataset confirmed the prevalence of the genera *Thalassiosira* and *Guinardia* during spring and summer and the relative higher contribution of *Navicula* species in winter, but gave a different picture of the seasonal succession within the diatoms since the metabarcoding approach allowed deciphering the annual sequence of a pool of persistently dominant nanodiatom taxa, such as the genera *Minidiscus*, *Cyclotella*, *Arcocellulus/Minutocellus* or the species *Thalassisira minima* (Fig 5C). In winter, *Minidiscus comicus* appeared as the dominant species while from April, and all along the summer and autumn, the contribution of *Thalassiosira* spp., *Cyclotella* and *Arcocellulus/Minutocellus* increased sequentially. While inter-annual variations were observed in the yearly sequence and contribution of dominant OTUs when individual years were considered (Fig. S6), the overall dominance of the smallest taxa was observed every year.

### 3.4 Temporal structure of planktonic protist community and ecological drivers

The use of a dbMEM analysis to decompose the temporal patterns of the community allowed us to detect and investigate the environmental and biological processes involved in the control of protist assemblages’ dynamics at different timescales (Fig. 6). The dbMEM eigenfunctions were retained by forward selection (47 and 52 positive MEMs, according to the morphological and metabarcoding datasets) and explained 48.9% of the species and 52.2% of the OTUs variability in community composition, respectively. As expected, seasonality - expressed in the first 2 constrained axes of the RDA - explained most of the observed temporal variability (RDA1: 19.8-17.8% and RDA2: 11.5% and 9.3%, for morphological and metabarcoding datasets respectively; Fig. 6B, D). For both datasets, the winter and summer assemblages on the one hand, and the autumn and spring assemblages on the other, were clearly distinguished on axes 1 and 2. Spring assemblages showed more interannual variability, especially when the morphological dataset was considered (Fig. 6A). The annual cycle was better delineated when the metabarcoding dataset was considered (Fig. 6C). For both datasets, the taxa/OTUs with the highest RDA1 and RDA2 scores corresponded to dominating species (section 3.3 and Fig. 5) and displayed clear seasonal variations in terms of cells or reads abundances (See Table 1 and Fig. 7A, B). For the morphological datasets, the pelagic chain forming *Guinardia delicatula* and *Thalassiosira levenderi/minima* and the benthic or tychopelagic taxa *Fragilaria/Brockmanniella*, *Paralia sulcata* and *Psammodictyon pandutiforme* had the highest scores for RDA1 and/or RDA1. For the metabarcoding dataset, the OTUs with the highest RDA1 and 2 scores also included *G. delicatula*, but pointed as well to nanoplanktonic diatoms (such as *Minidiscus comicus*), and to species belonging to other phyla or classes such as the Dinophyceae and Cercozoa, all displaying strong seasonality (Fig. 7B and Table 1).

**Figure 6.**
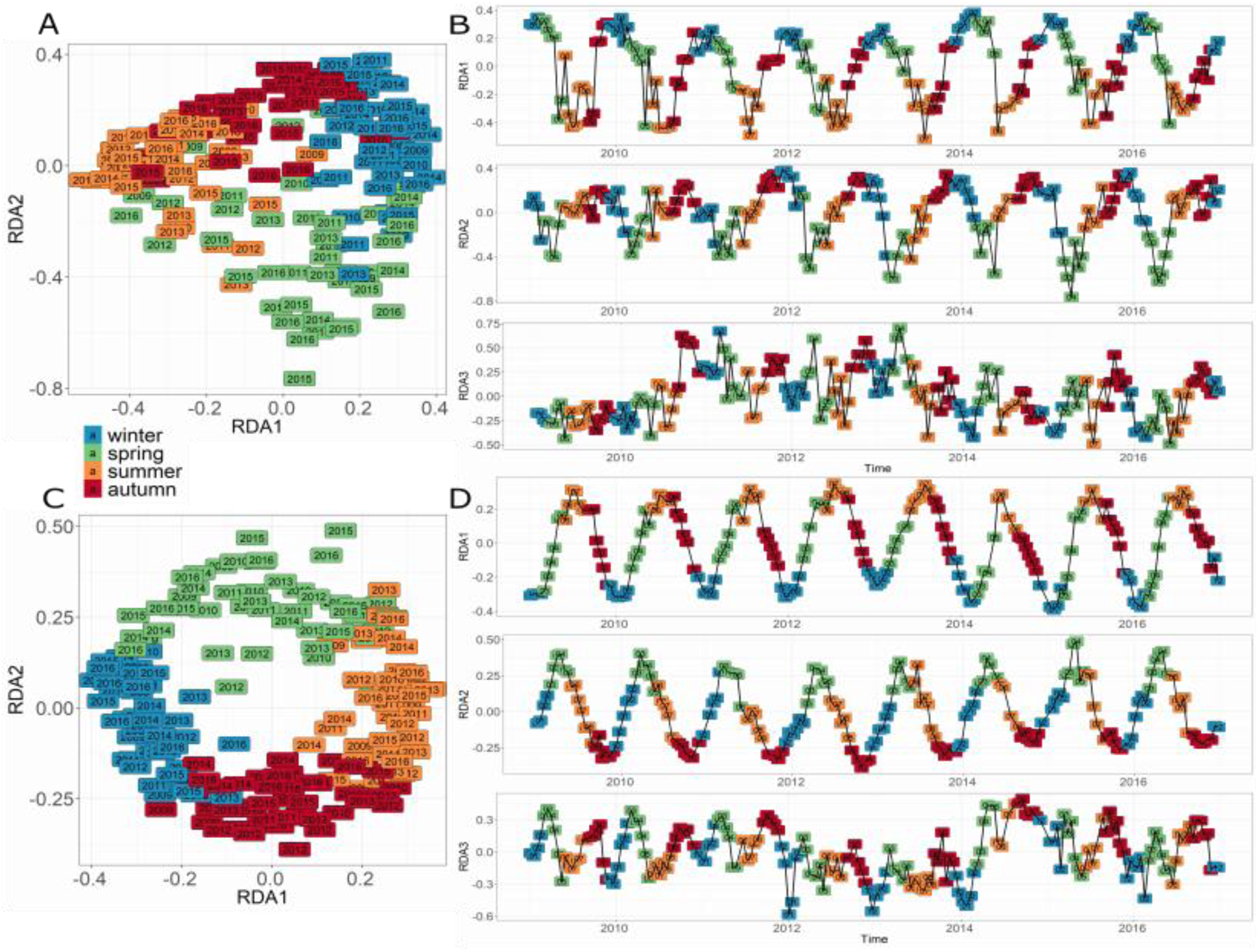
Similarity of protist communities (RDA analysis) in monthly samples over the period 2009-2016 at SOMLIT-Astan sampling station for morphological microscopy. **(A**, **B), and DNA metabarcoding (C**, **D) datasets.** (**A**, **C**): Annual cycle of protist communities obtained by ordination of the monthly samples through a redundancy analysis (RDA) explaining (**A**) 48,9 % and (**B**) 52,2 % of the total variance of the community, respectively. (**B**, **D**): Decomposition of RDA axes that reveals seasonal pattern (RDA1; 19,8-17,8 % and RDA2; 11,5-9,3 %) and biannual broad scale oscillation (RDA3; 4,8-3,9 %).

**Figure 7.**
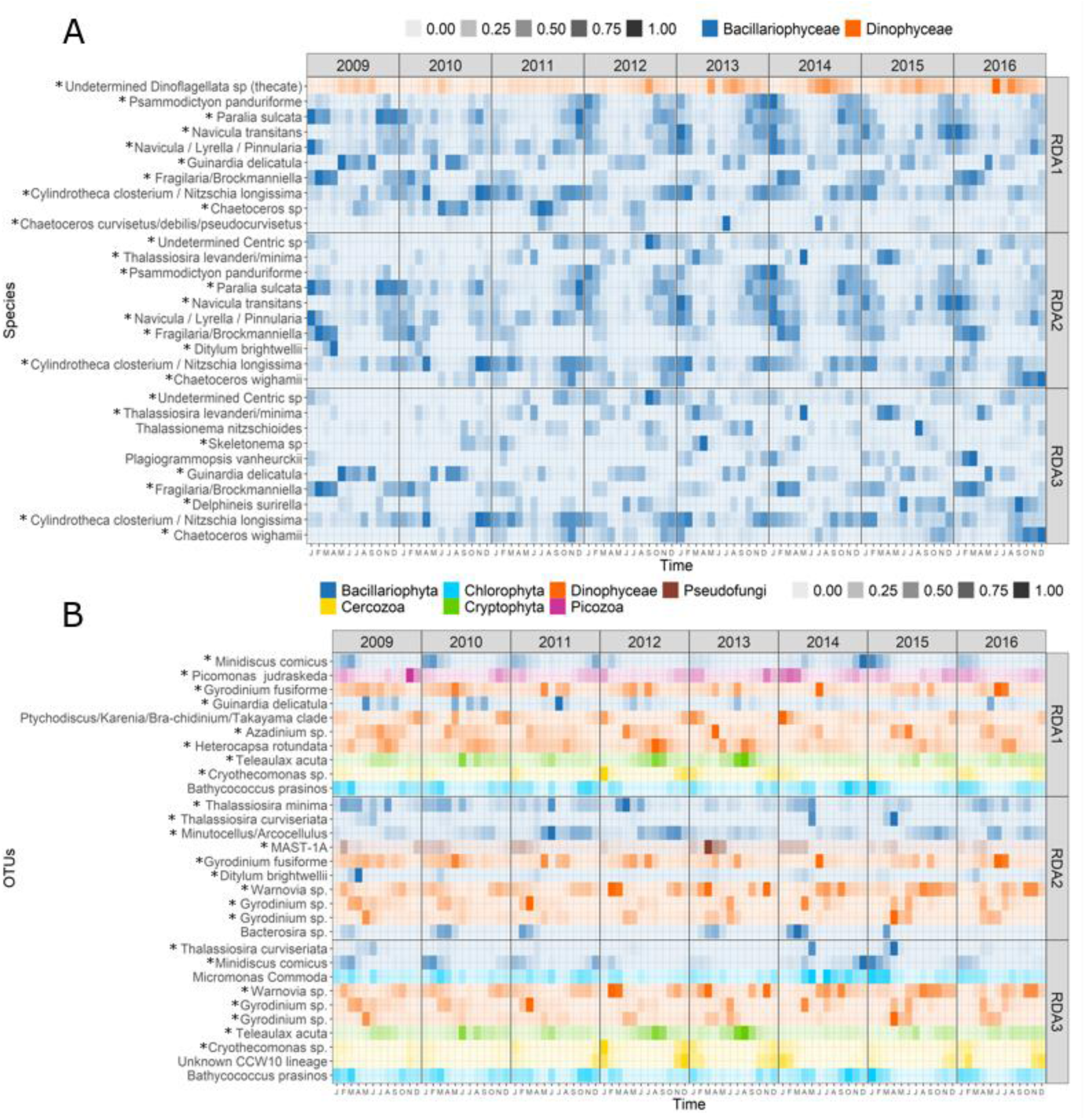
Monthly mean abundance (2009-2016), at the SOMLIT-Astan sampling site, for (A) the morphological species and (B) molecular OTUs as a function of the first three RDA axes (see Figure 6). For each RDA axis the (**A**) 10 species and (**B**) 10 OTUs with the highest score were selected. The * indicates dominant OTUs (reported in Fig.5).

**Table 1.** Species and OTUs driving the seasonal oscillation showed in the RDA axes 1 and 2 and the biannual broad scale oscillations observed in axes 3. The 10 species/OTUs with the highest scores in each relative axes of the RDA were selected. The scores and the relative contributions to total abundance of the resulting list of species/OTUs are shown. For metabarcoding data, see Material and methods section and table S1 for assignations details.

| Species / Assignment | Contribution to variation axis 1 (%) | Contribution to variation axis 2 (%) | Contribution to variation axis 3 (%) | Contribution to total abundance (%) |
| --- | --- | --- | --- | --- |
| **Morphological dataset** |  |  |  |  |
| *Chaetoceros curvisetus/debilis/pseudocurvisetus* | 2.4 | - | - | 1.3 |
| *Chaetoceros* sp. | 3.0 | - | - | 2.7 |
| *Chaetoceros wighamii* |  | 3.7 | 18.3 | 3.0 |
| *Cylindrotheca closterium* / *Nitzschia longissima* | 3.9 | 2.0 | 2.0 | 5.9 |
| *Delphineis surirella* | - | - | 0.8 | 1.6 |
| *Ditylum brightwellii* | - | 1.6 | - | 0.5 |
| *Fragilaria/Brockmanniella* | 20.2 | 8.5 | 10.5 | 5.7 |
| *Guinardia delicatula* | 28.2 | - | 8.48 | 8.7 |
| *Navicula* / *Lyrella* / *Pinnularia* | 2.5 | 1.33 | - | 2.7 |
| *Navicula transitans* | 2.4 | 3.0 | - | 2.7 |
| *Paralia sulcata* | 12.5 | 15.7 | - | 10.6 |
| *Plagiogrammopsis vanheurckii* | - | - | 0.97 | 0.4 |
| *Psammodictyon panduriforme* | 1.9 | 4.69 | - | 1.8 |
| *Skeletonema* sp. | - | - | 43.2 | 4.0 |
| *Thalassiosira levanderi/minima* | - | 45.7 | 5.0 | 6.6 |
| *Thalassionema nitzschioides* | - | - | 0.91 | 0.7 |
| *Undetermined Centric* | - | 2.5 | 1.2 | 1.9 |
| *Undetermined Dinoflagellata (thecate)* | 2.9 | - | - | 6.2 |
| **Metabarcoding dataset** |  |  |  |  |
| *Bacterosira* sp. | - | 1.9 | - | 0.3 |
| *Bathycoccus prosinos* | 1.5 | - | 1.86 | 0.9 |
| Unknown CCW10 lineage | - | - | 2.3 | 0.3 |
| *Cryothecomons* sp. | 3.2 | - | 6.4 | 1.4 |
| *Teleaulax acuta* | 2.4 | - | 2.2 | 2.2 |
| *Heterocapsa rotundata* | 2.1 | - | - | 2.4 |
| *Gyrodinium* sp. | - | 5.7 | 2.9 | 2.2 |
| *Azadinium* sp. | 2.2 | - | - | 1.1 |
| *Gyrodinium* sp. | - | 7.1 | 2.5 | 1.7 |
| *Ptychodiscus/Karenia/brachydinium/takayama clade* | 1.4 | - | - | 0.5 |
| *Warnovia* sp. | - | 2.2 | 7.6 | 3.2 |
| *Ditylum brightwellii* | - | 2.2 | - | 0.4 |
| *Guinardia delicatula* | 4.1 | - | - | 1.5 |
| *Gyrodinium cf fusiforme* | 1.8 | 2.2 | - | 2.1 |
| MAST-1A | - | 2.3 | - | 0.5 |
| *Micromonas commoda* | - | - | 1.9 | 0.8 |
| *Picomonas judraskeda* | 1.5 | - | - | 0.7 |
| *Minidiscus comicus* | 10.0 | - | 6.1 | 2.7 |
| *Minutocellus/Arcocellulus* | - | 2.2 | - | 1.0 |
| *Thalassisira minima* | - | 2.0 | - | 1.2 |
| *Thalassisira curviseriata* | - | 3.0 | 2.7 | 0.7 |

The axis 3 of the RDA (4.8 and 3.9 % of the variance explained for the morphological and metabarcoding datasets respectively, Fig. 6B, D) expressed broad scale oscillations and a persistent biannual rhythm in the protist community dynamics. In the morphological dataset, *Skeletonema* sp. contributed most to axis 3 of the RDA. *G. delicatula* and *Chaetoceros wighamii* also showed high contribution. In the metabarcoding dataset, the winter diatom *M. comicus* and the Cercozoan *Cryothecomonas*, that exhibit a parasitic life styles, had the highest contribution to this axis. Our analyses showed that some of the OTUs were not detectable every year, suggesting an influence of the environment on the temporal patterns of some protists.

To investigate the environmental factors that primarily drive seasonal protist assemblages, we calculated Spearman rank correlation coefficients between the potential explanatory variables and the first 3 axes of the RDA (Fig. 8A, B). Here, we considered the environmental variables selected by forward selection for both datasets, namely: temperature, phosphates (PO_4_^3-^), silicates (SiO_4_^-^), ammonia (NH_4_^+^), Chl-a, salinity, Suspended Matter (MES) and the North Atlantic Oscillation (NAO) Index. Oxygen was selected only for microscopy, and PAR, nitrate (NO_2_^-^), pH, and Delta N15 for metabarcoding. The analyses suggested that macronutrients (PO_4_^3-^, NH_4_ and to a less extent SiO_4_ ), together with temperature, PAR, and Chlorophyll-*a* showed the highest correlations with RDA1. Temperature, salinity, oxygen, pH and NO_2_^-^ showed the highest correlations with RDA2 (Fig. 8A, B). Even though environmental variables alone only accounted for 5% of the variance, a large part of the variations in the community was explained by the temporal structure of the environmental factors (26% and 24% for both datasets, respectively, Fig. 8C, D). The temporal organization of the community explained most of the variations when considered together with the environment (47% and 49%). These results suggest that the temporal structures of the environmental factors are important drivers of the annual succession of species, along with instrinsic community effects such as species interactions, reproductive dynamics, and stochastic events not considered here.

**Figure 8.**
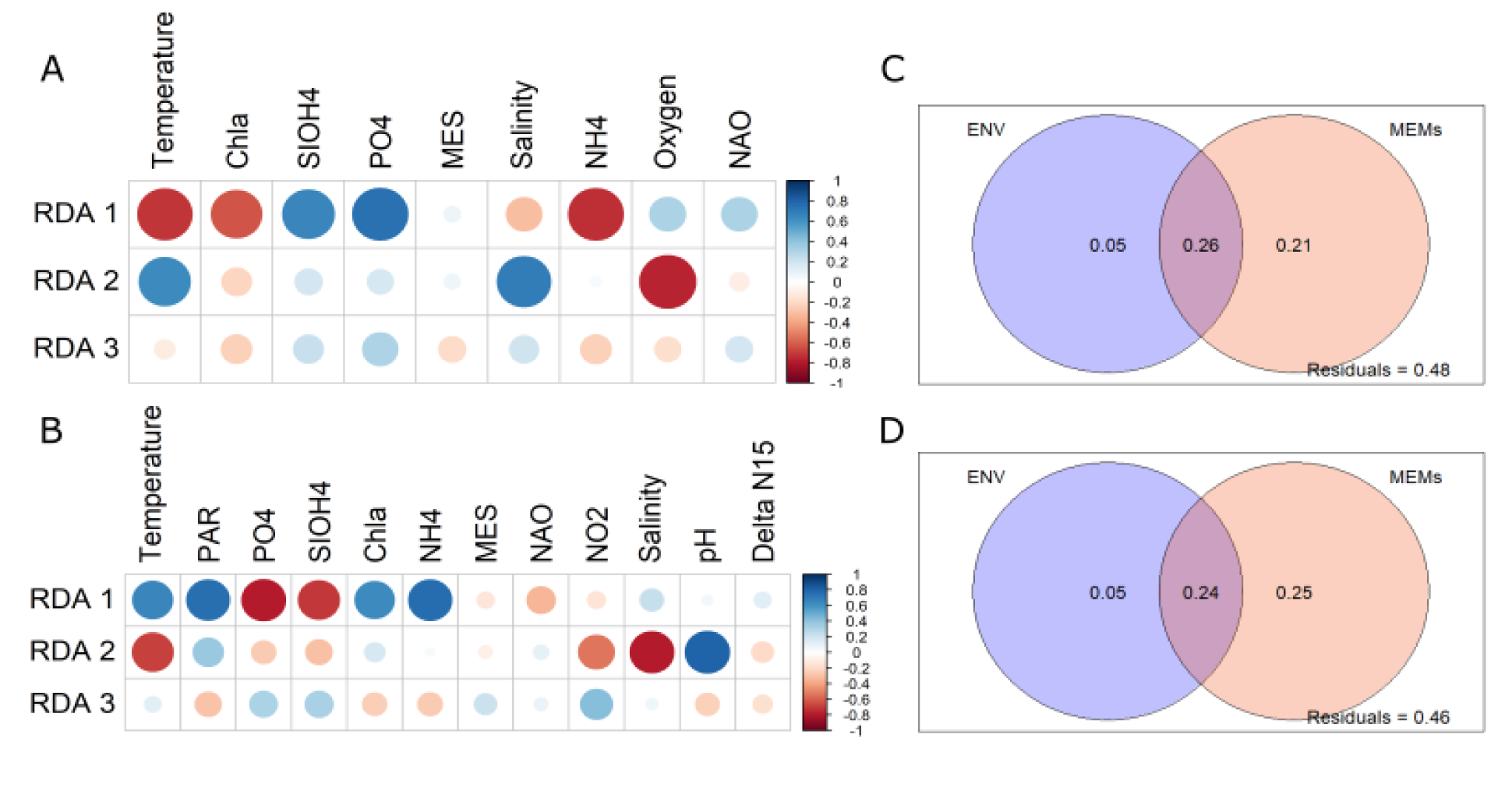
Spearman correlation calculated between the environmental variables and the RDA axes (A, C), and variance partitioning analyses between environmental drivers and dbMEM (B, D). Spearman correlations were computed between each axes of the RDA and each environmental parameter selected for (**A**) morphology and (**B**) metabarcoding. Variance partitioning between selected environmental variables and dbMEM was also calculated for (**C**) morphology and (**D**) metabarcoding data, respectively.

## 4. Discussion

### 4.1 Predictable cyclicity of protists successions in coastal pelagic habitats

Annual successions of planktonic protists, in particular phytoplankton, have been observed by early planktonologists (for diatoms and dinoflagellates, see for example Allen, 1936; Gran and Braarud, 1935) and have inspired the founding theories of ecological successions (Margalef, 1963; Margalef et al., 1958, 1978). Recent temporal analyses of DNA metabarcoding datasets from a wide range of biogeographical regions have confirmed that such patterns are a general common feature of marine microbes, including prokaryotes and all protists (Marquardt et al., 2016; Egge et al., 2015; Piredda et al., 2017). Some also noted the overall stability of the yearly reoccurring sequences of taxa at the decade scale (see Fuhrman et al., 2015 for bacteria and Lambert et al., 2019 or Giner et al., 2019 for protists).

Using morphological and DNA metabarcoding approaches, we clearly identified annual succession patterns of taxa in the Western English Channel over the period 2009-2016. The cyclic pattern was more distinct using metabarcodes (see Fig. 6A *versus* 6C), however: by including some of the smallest eukaryotes, a greater diversity was therefore assessed. Compared to microscopic counts (146 morphological taxa of mainly microphytoplankton groups), genetic sequences data enabled to reach a much finer taxonomic resolution (15,271 OTUs belonging to 53 different phyla and classes) and gave access to the dynamics of nano-planktonic taxa (including some pico-) that are dominant at our site. For example, the pico-planktonic green algae *Ostreococcus lucimarinus*, *Bathycoccus prasinos* and *Micromonas* spp. or the nanoplanktonic diatoms *Minidiscus variabilis* and *M. comicus*, are known major players of the microbial communities in the Western English Channel (Not et al., 2004; Foulon et al., 2008; Arsenieff et al., 2020). DNA metabarcoding could also capture the dynamics of naked dinoflagellates taxa (*Gyrodinium* and *Gymnodinium* species) and heterotrophic, parasitic or endosymbiotic microeukaryotes such as the MAST, *Cryothecomonas*, and Syndiniales species. Taxa such as *Cryothecomonas* that infects diatoms and especially the genus *Guinardia* (Drebes et al., 1996; Peacock et al., 2014), and the Syndiniales that parasite dinoflagellates (Chambouvet et al., 2008) are involved in the control of phytoplankton blooms and thus in the overall stability of the system. These taxa, which cannot be easily monitored using microscopy, are prominent all year round in terms of DNA read abundances, and contributed significantly to the seasonal temporal variations captured by the two first axes of our RDA analyses. Marine benthic protists indeed show high and distinctively different diversity compared to planktonic species (Patterson et al., 1989), especially in the Cercozoa (Forster et al., 2016), one of the dominant phyla at SOMLIT-Astan.

Such regularities and stability in community composition over 8 years in the surface waters of this megatidal region could be related to different factors. By transporting planktonic species from and to adjacent habitats, tidal currents are known to increase plankton dispersal, which is an important process in structuring communities (Vellend et al., 2010). Therefore, tidal amplitude is intensifying the exchanges between communities associated to the diverse habitats of the Morlaix Bay (Cabioch et al., 1968; Dauvin et al., 2008; Gac et al., 2020). In habitats influenced by tidal mixing, high contributions of benthic protists to the water column communities are classically observed (Hernandez-Farinas et al., 2017); this phenomenon is amplified in winter, when pelagic species are less abundant and winds increase vertical mixing (Mann & Lazier, 1991). Nevertheless, the induced high rates of emigration and immigration do not apparently disrupt the seasonal oscillations in diversity that was captured by the first 2 axes of our RDA analyses (Fig.6). By increasing the overall diversity and by enhancing bentho-pelagic coupling (and potentially the interactions between species), these forces may favor the overall stability of the system (Cardinale et al., 2012). Our dbMEM analyses allowed us to uncover a clear and persistent biannual rhythm (axis 3 of our RDA) in the protist community dynamics. This biannual pattern was also observed in the monthly turnover rates of species over the 8 years (Fig. S4). However, no clear relationship was found with annual variation in tidal amplitude which was not among the environmental factors selected for correlation analyses (Fig.8).

We found a strong correlation between the first two axes of the RDA and temperature, macronutrients and light (Fig. 8), suggesting that physical factors related to the annual climate cycle are imposing synchrony to the protist seasonal dynamics (Cloern, 1996; Sommer et al., 2012). Nevertheless, our results showed a more holistic vision, where most part of the community dynamics was explained by the temporal structure of both environmental and intrinsic factors (Fig. 8B and D).

### 4.2 The annual sequence of dominant protists in temperate tidally-mixed habitats

With observations conducted over 8 years using both microscopy and DNA metabarcoding, our study improves our knowledge of pelagic protists in a tidally-mixed coastal environment. In terms of phytoplankton, our study confirmed the importance of diatoms (by far the most numerous taxa >10µm enumerated under microscopy), dinoflagellates and green algae, but also highlighted the importance of Cryptophyta. The DNA metabarcoding analysis provided new data about the seasonal sequence of important heterotrophic dinoflagellates (i.e., Dinophyceae and Syndiniales) harboring diverse trophic modes, and that of parasitic Cercozoa. Along the years 2009-2016, the prominence of the chain-forming species *Guinardia delicatula* during spring and summer was confirmed by both the morphological and metabarcoding datasets. This species is emblematic of the spring and summer diatoms bloom in the Roscoff area (Grall, 1972; Martin-Jezequel, 1992; Sournia et al., 1995; Guilloux et al., 2013; Arsenieff et al., 2019). It is one of the most recognized species of planktonic communities in the English Channel and North Sea (Widdicombe et al. 2010; Caracciolo et al., 2021) and appear to be particularly successful in temperate tidally-mixed habitats (Gomez and Souissi, 2007; Wiltshire et al., 2008; Peacock et al., 2012; Schlüter et al., 2012; Hernandez-Farinas et al., 2014). Our analyses also tracked the classical annually repeated sequence of diatoms that involves the development of microplanktonic pelagic chain-forming species in spring (typically *Thalassiosira* spp., *G. delicatula*, *Chaetoceros* spp.), as well as benthic and tychopelagic species in winter (e.g. *Paralia* sp. and *Navicula* spp.). An annual sequence of nanodiatoms (involving species of the genera *Minidiscus*, *Thalassiosira* and *Arcocellulus / Minutocellus*) was specifically revealed by the metabarcoding approach. The prevalence of nanodiatoms, and especially of *Minidiscus* spp. at the SOMLIT-Astan station has actually been confirmed by cultural approaches from the SOMLIT-Astan station (Arsenieff et al., 2020). Nanodiatoms are have been identified as prominent members of diatoms assemblages in other marine systems when adequate detection techniques (cultural approaches, electron microscopy or HTS) were implemented (Leblanc et al. 2018; Ribera l’Alcalà et al. 2004; Percopo et al. 2011).

If microphytoplanktonic dinoflagellates are usually present at relatively low abundances in species microscopic counts in tidally-mixed waters off Roscoff (Sournia et al., 1987; Guilloux et al., 2013), the contribution of dinoflagellates reads in the molecular dataset was high all year round according to this study. Sequences corresponding to the dominant reads were mostly assigned to nanoplanktonic species or naked species. Two OTUs assigned to the genus *Heterocapsa* including the thecate species *H. rotundata* dominated read counts for the whole period. This ubiquitous mixotrophic dinoflagellate, that has the potential to switch from phototrophy to partial heterotrophy (Millette et al., 2017), may be favored at our tidally-mixed coastal site, especially in August when light starts decreasing. Interestingly, *H. rotundata* was also identified as a dominant taxon in the adjacent Penzé estuary (Chambouvet et al., 2008), and as most abundant in recent microscopic counts obtained from our time-series station where nanoplanktonic dinoflagellates were targeted (data not shown). Some other dominant dinoflagellate OTUs detected in our metabarcoding dataset are either heterotrophic or potentially mixotrophic (*Gyrodinium, Gymnodinium, Azadinium, Warnovia* etc.), and some of them are purely parasitic (Syndiniales). The naked dinoflagellates *Gyrodinium* and *Gymnodinium* spp. were also identified as prominent members of the phytoplankton community in the stratified waters of the WEC, off Plymouth (Widdicombe et al., 2010) and showed an increasing trend in abundance after 2001 (Hernández-Fariñas et al., 2014). These dinoflagellates, that seem to thrive all year round, may be key predators for diatoms. The increasing trend in average abundance of some dinoflagellates and the decrease in diatoms has been recently documented in the Central North Atlantic Ocean and in the North Sea (Leterme et el., 2005; Zhai et al., 2013), as well as in the English Channel (Widdicombe et al., 2010).

Cryptophyta are important members of protists communities in coastal waters. Their prominence in different regions of the ocean has been revealed using microscopy (Jochem, 1990), but also via flow cytometry, since the phototrophic members of this group can be distinguished based on its phycoerythrin fluorescence (Dickie, 2001). Recent DNA metabarcoding analyses have also revealed their prominence in coastal waters at Helgoland Roads, North Sea (Käse et al., 2020). At the SOMLIT-Astan station, sequences identical to different species of the genus *Teleaulax* were abundant in read counts. The highest proportion of Cryptophyta reads were assigned to *Plagioselmis prolonga* (=*Teleaulax amphioxeia*), a phototrophic species with a bentho-pelagic life-cycle (Altenburger et al., 2020) involved in complex symbioses with the ciliate *Mesodinium rubrum* (Qiu et al., 2016). Besides, the later species has an interesting behavior consisting of periodic dispersion away from the strong superficial tidal currents, thus minimizing flushing losses (Crawford and Purdie, 1992).

We are aware that the description of the typical seasonal sequence of protists species provided herein is still incomplete. Both microscopy and metabarcoding can provide biased data, since the former does not consider the smallest taxa while the later can overestimate or underestimate the proportions of taxa for which DNA is more easily extracted and amplified (Santi et al. 2021). For example, the contribution of dinoflagellate to sequences reads obtained from natural samples is a commonly reported bias due to the very high number of 18S copies in dinoflagellates (Gong and Marchetti, 2019). In our study, the contribution of Haptophyta was probably underestimated since most species in this group are nano- or picoplanktonic and were thus not reported in our morphological dataset, as the primers used do not perfectly match with sequences of this group (McNichol et al., 2021). However, the two datasets we used are complementary and allowed us to add important information about the dynamics of dominant protists thriving in permanently mixed waters of the Western English Channel. A deeper analysis of species dynamics in the different phyla for the metabarcoding dataset will certainly provide more information in the future, especially since reference sequences databases and taxonomic frameworks (required for accurate assignations to genus or species levels) are constantly being updated and curated (Guillou et al., 2013; Berney et al., 2017; Glöckner et al., 2017; del Campo et al., 2018).

### 4.3 Environmental *versus* community intrinsic drivers of protistan plankton seasonal dynamics

Plankton phenology and seasonal species successions are periodic processes that are tightly phased to astronomical forcing and associated annual cycles of temperature, solar radiation, atmospheric heat input and photoperiod (Sverdrup, 1953; Cushing, 1959, Winder & Cloern, 2010). While being structured by local physical and chemical conditions and biological self-organization process, microbial eukaryotic communities, as a feed-back, strongly impact biogeochemical cycles (Arrigo, 2005; Falkowski et al., 2008; Fuhrman, 2009). Spearman correlation coefficients calculated between environmental variables and the first three axes of the RDA, confirmed a strong phasing between seasonal variations of environmental parameters and protists assemblages. According to our analyses, the resilient seasonal pattern in community dynamics is tightly linked to the temporal structure of Photosinthetically Active Radiation, temperature, salinity and macronutrients (PO_4_^3^, SiO_4_^2-^ , NH_4_^+^, Fig. 8A, and in a less extent NO_2_^-^, Fig. 8B). Thus, we identified the earth tilt and rotation rhythms determining light intensity as the primary driver of the observed plankton dynamics, in association with cyclical variations in water temperature and principal macronutrients concentration. Those factors are more important with respect to local hydrodynamic conditions, nevertheless, local factors such as salinity and pH showed high correlation with the second axes (Fig. 8B), suggesting that phytoplankton variability also depends on additional factors in nearshore waters, such as tides and river runoff (Cloern, 1996). The presence of Delta N15 in the selected environmental variables (Fig. 8B), suggests that agriculture, intensively practiced in our region of interest, could also influence the community structure that we observe at different time of the year.

The large contribution of autocorrelation to the variance of the community (Fig. 8C) suggests that intrinsic biological factors (i.e., species interactions, reproductive dynamics, and/or self-regulation of species development; in other words, self-organization properties of the whole biological community; Odum, 1988; Picoche and Barraquand, 2019) are also critical, and significantly contribute to pacing the plankton community. Biotic factors, such as species interactions, partly explain the striking resilience in species turnover also observed for bacterioplankton along a decade (Fuhrman et al., 2015). Microscopic organisms are indeed known to be involved in complex and dynamical networks of interactions (i.e., grazing, parasitism, mutualism, quorum sensing, etc; Kivi et al., 1993; Dakos et al., 2009; Platt et al., 2009; Bjorbækmo et al., 2020) that are tightly regulating the dynamics of individual species within the whole community structure. Recently, in the WEC at L4 station predator-prey interactions were investigated supporting our hypothesis that they play a role in influencing temporal changes in plankton populations (Barton et al., 2020).

Bi-annual variations in the protist community dynamics were also identified from analyses of both dataset (Fig. S4 and Fig. 6). In some ecosystems, rhythmic depletions of resources appear to be at the origin of bimodality (and multimodality) in phytoplankton dynamics (Mellard et al., 2019), however, in our tidally-mixed coastal station, nutrients are never completely depleted (Fig. 2). The bi-annual rhythm could be triggered by the yearly variations of the tidal cycles that are probably at the origin of cycles of enhanced bentho-pelagic coupling; and recently, a reorganization of marine food webs due to strong pelagic-benthic coupling in coastal areas has been reported (Kopp et al., 2015). The number of benthic species detected as prominent in surface waters suggests a tight coupling between benthic and pelagic compartments in the English Channel, which is strengthened in winter, when tidal mixing or winds provoke the resuspension of sediments in the water column. However, in our dbMEM analysis, neither tidal amplitude nor wind appear as a major influential parameter. This yearly bimodality could then be caused by intrinsic plankton biological factors such as endogenous rhythmicity or interactions between species. To better decipher how intrinsic biotic interactions could drive the dynamics of these communities, modelling approaches that take into account biotic interactions (e.g., Picoche and Barraquand, 2019) should be explored, integrating the whole taxonomic and functional spectrum that coexist in space and time, including viruses, prokaryotes, and metazoans.

## 5. Conclusions and perspectives

This study describes the seasonal dynamics of protist communities in a temperate coastal site, complementing early studies (Cushing, 1959; Margalef, 1963; Sommer et al., 1986; Widdicombe et al., 2010) from the same site which considered only diatoms and dinoflagellates. It appears that seasonal successions of protists are primarily paced by astronomical rhythms and may be directly influenced by the resulting temporal variations of the physical and biogeochemical parameters (i.e., PAR, temperature, nutrients, salinity). Intrinsic, plankton self-organization processes are also involved in these annual oscillations. In environments such as the coastal waters of the EC that support one of the busiest shipping lanes in the world, important fishing ports, and an increasing demographic pressure, these seasonal cycles may actually be particularly vulnerable to the combined effects of natural climate variability and local anthropogenic forcing (Dauvin et al. 2008, Tréguer et al. 2014, Gac et al. 2019, Siano 2021). In this context, monitoring activities involving both classical microscopy and metagenomics approaches, such as those conducted along the EC coasts (Breton et al., 2000; Widdicombe et al. 2010; Hernandez-Farinas et al., 2017; Kenitz et al. 2017; Käse et al., 2020) should be maintained and developed on the long term. These longitudinal surveys are critical to track and predict future changes that may disrupt the overall resilience of the system, in order to ultimately identify and deploy protective measures to guarantee all the services that these systems provide to the society (Cardinale et al., 2012).

## Acknowledgements

The authors would like to thank the captains and crew of the Neomysis research ship for their help during sampling at the SOMLIT-Astan station. We are also grateful to the RCC for the maintenance of phytoplankton strains isolated from this station that served for the assignation of some of the genetic sequences. We also thank Cédric Berney and Benjamin Alric for stimulating discussions, and for suggestions to improve the manuscript. Daniel Vaulot is acknowledged for his work on the phytoplankton counts databases. This study was supported by a PhD fellowship from Sorbonne University to MC, the French government research agency programs CALYPSO (ANR-15-CE01-0009), BIOMARKS (“Investissement d’avenir”, ANR-08-BDVA-0003) and OCEANOMICS (ANR-11-BTBR-0008), the CNRS-INSU EC2CO CYCLOBS research project grant and the Gordon and Betty Moore Foundation through the UniEuk grant GBMF5275.

## Authors contributions

NS, FN, NH and MC designed the study. FRJ, TC and the crew of the Neomysis sampled onboard. FRJ and LG produced the taxa counts. MH helped with the construction and maintenance of the phytoplankton counts databases. FRJ, SF and NS contributed to the corrections and validation of the taxonomic counts dataset. FRJ and SR produced the genetic data. TC produced the hydrological data and TC and YB contributed to the validation of the hydrological data validation. JPG produced the final estimations of pH and FCO2. EG provided the map. SC helped with the calculations of the PAR data. MC and NH analysed the data and produced the scripts and final graphs. MC, NH, ET, FRJ, SR and NS wrote the manuscript. All authors contributed to the discussions that led to the final manuscript, revised it and approved the final version.

## Data accessibility

− Raw environmental data: https://meteofrance.com/, https://www.somlit.fr/en/, https://www.somlit.fr/en/, https://data.shom.fr/, https://modis.gsfc.nasa.gov/data/dataprod/, and https://coastwatch.pfeg.noaa.gov/
− Taxonomic counts input file : https://doi.org/10.5281/zenodo.5033180
− Metabarcoding raw data: It will be made available after acceptance of the manuscript.
− Metabarcoding input file: https://doi.org/10.5281/zenodo.5032451

