## Supplementary figures and tables for "Seasonal temporal dynamics of marine protists communities in tidally mixed coastal waters"

**Figure S1**

**Summary of analyses performed on the OTU contingency table.** Each white box represents a manipulation of the data necessary to arrive at the final result in the color box.

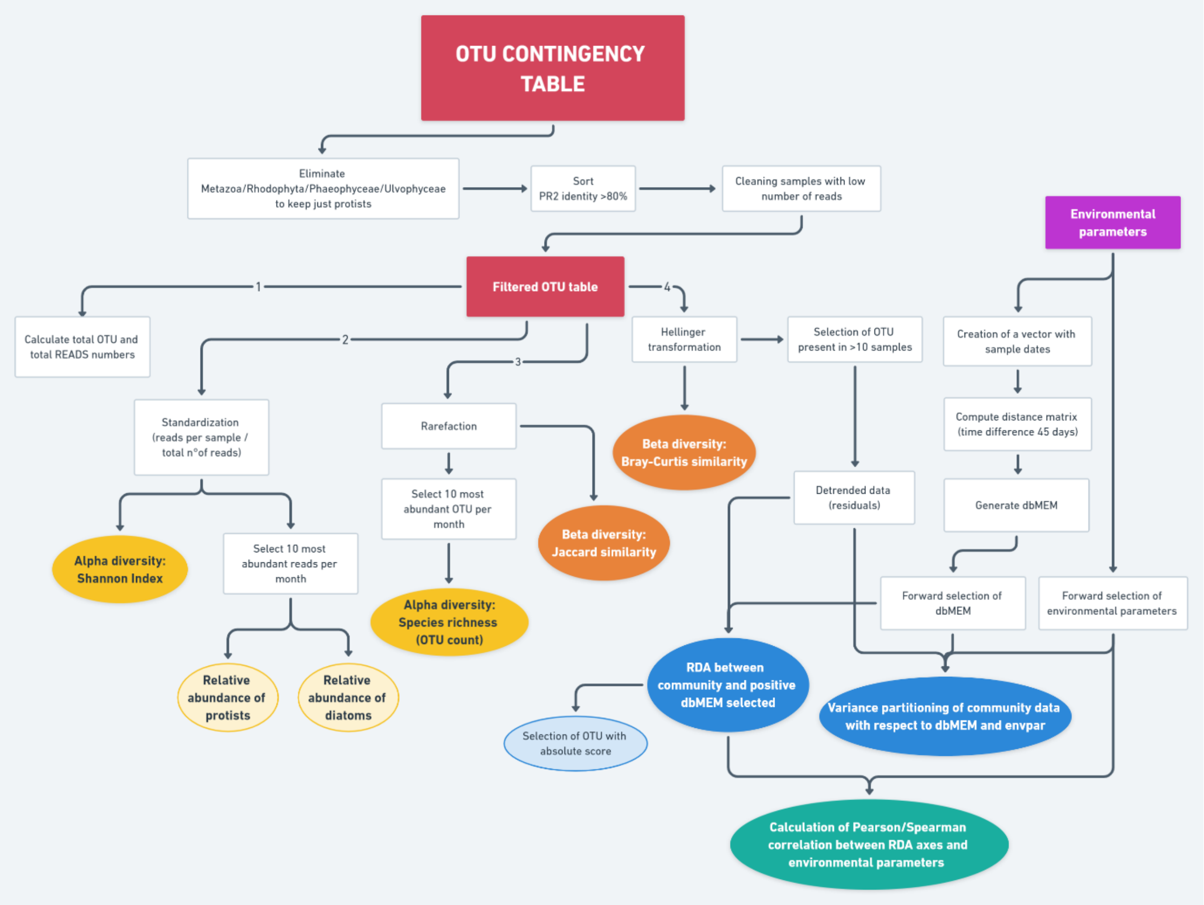

**Figure S2**

**Mean yearly variations recorded for the hydrological and meteorological parameters at the SOMLIT-Astan time-series station in the period 2009-2016.** All measurements were obtained for high neap tides periods. PAR8day is the photosynthetically available radiation calculated as the average light received during the 8 days that preceded each sampling dates. Kd490 is intended as the diffuse attenuation coefficient for downwelling irradiance at 490 nm (for more details about each parameter see Material & Methods section).

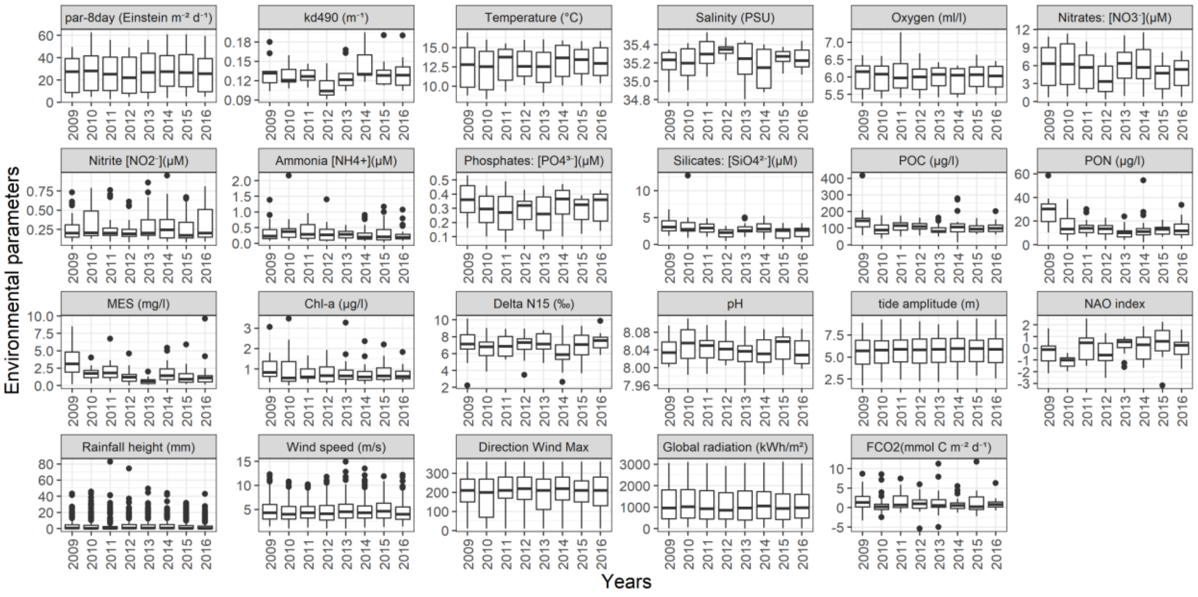

**Figure S3**

**Monthly variations of the Shannon Index.** For metabarcoding, alpha diversity was calculated at the class level or phylum level. The absence of data during some months for certain classes is linked to the absence of that OTUs during those months.

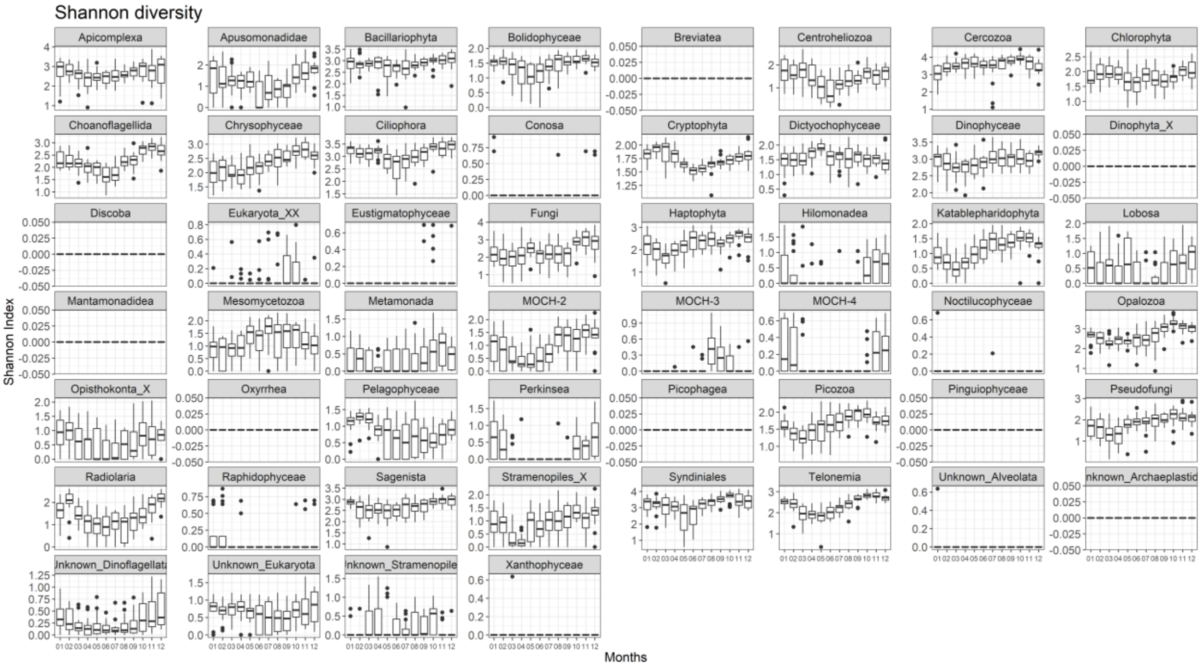

**Figure S4**

**Monthly variations in the ecosystem turn-over at the SOMLIT-Astan station for the period 2009-2016** as estimated from the protist community: (**A)** Bray-Curtis dissimilarities, (**B**) Jaccard distances as calculated from metabarcoding data and (**C**) monthly means euclidian distances as calculated from environmental data.

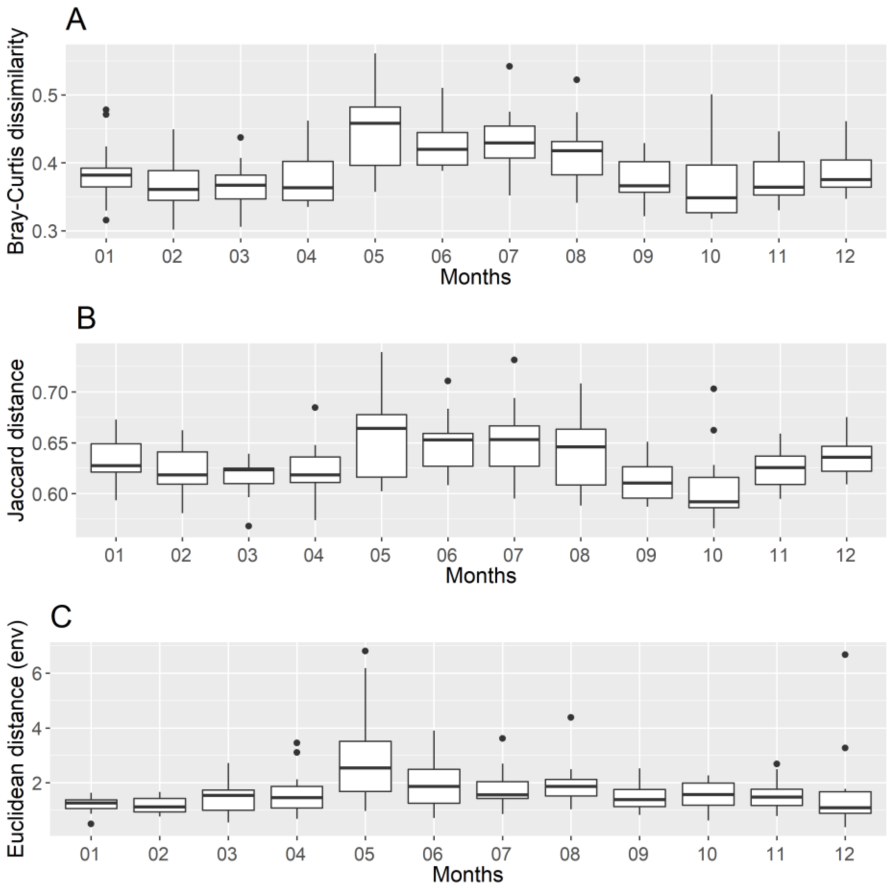

**Figure S5**

**Monthly variations in the cell abundance (A) and contribution to reads abundances (B) of dominating high-rank taxonomic groups at the SOMLIT-Astan time-series station over the period 2009-2016.**

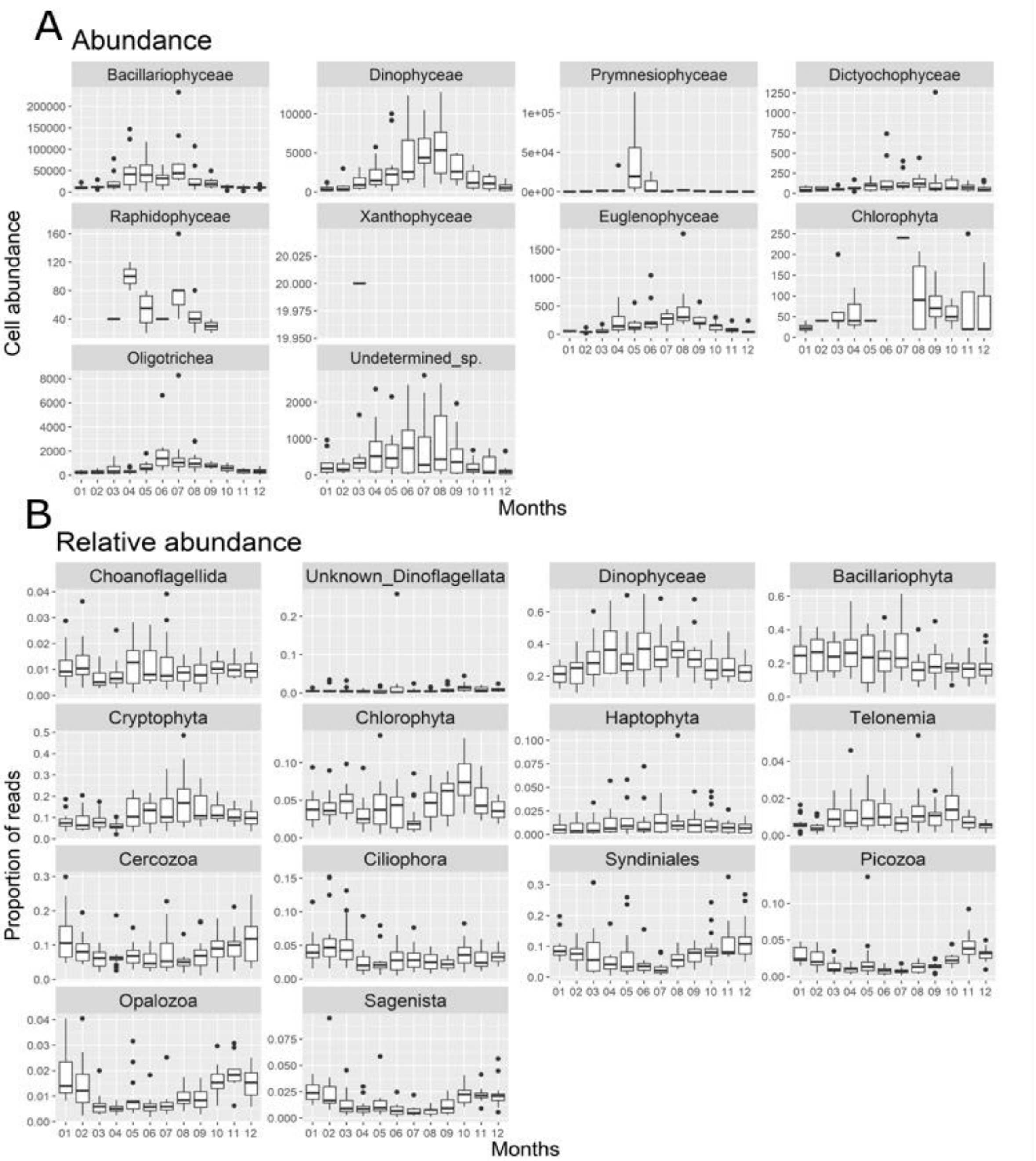

**Figure S6**

**Temporal variations in the monthly contributions of dominating protists or diatoms at the SOMLIT-Astan time-series station over the period 2009-2016.** (**A**) The contributions to total DNA reads abundance of the dominating OTUs; (**B**) contributions of the main diatoms to total species abundances; (**C**) contribution of the main diatoms to total diatom reads abundances. OTUs/species selected were the 10 most abundant for at least one month, when mean monthly abundances were taken into account (5 most abundant for diatoms).

**A**

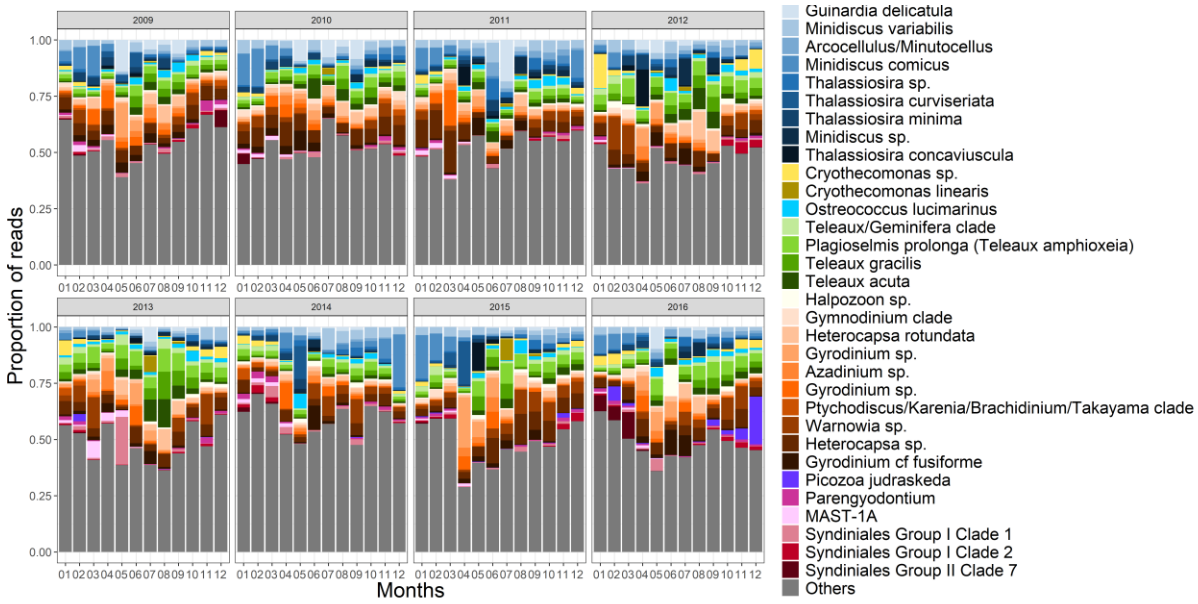


**B**

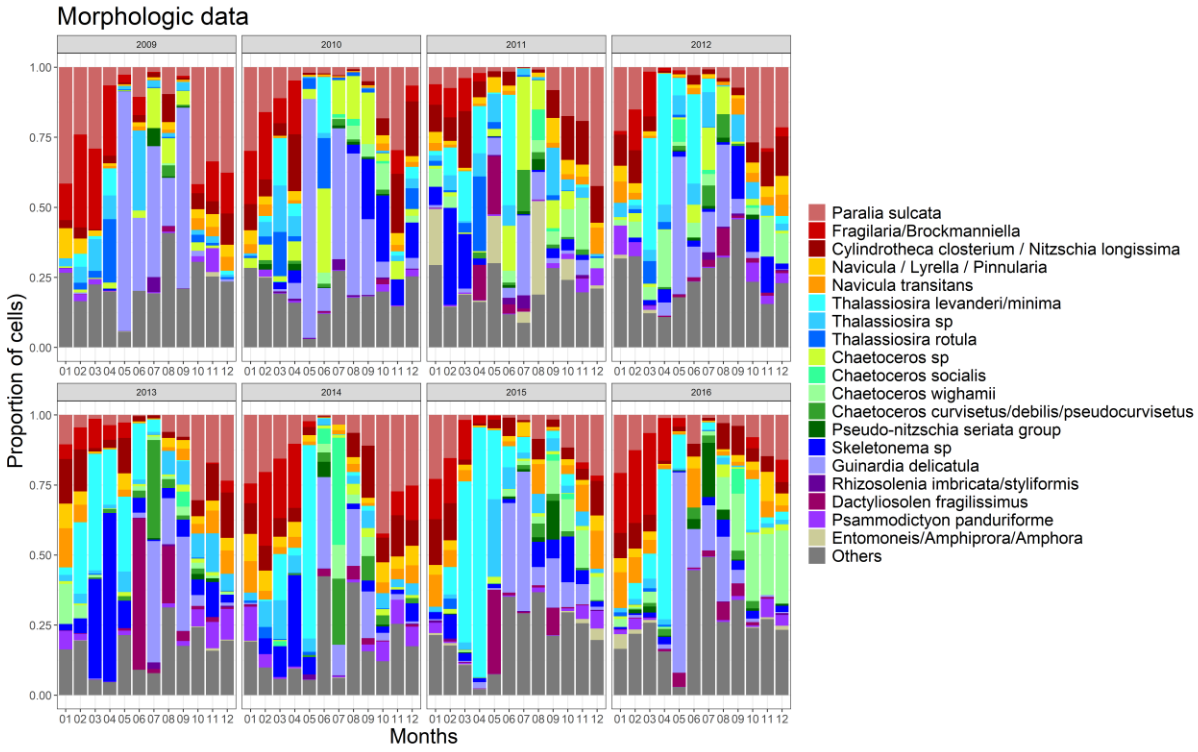

**C**

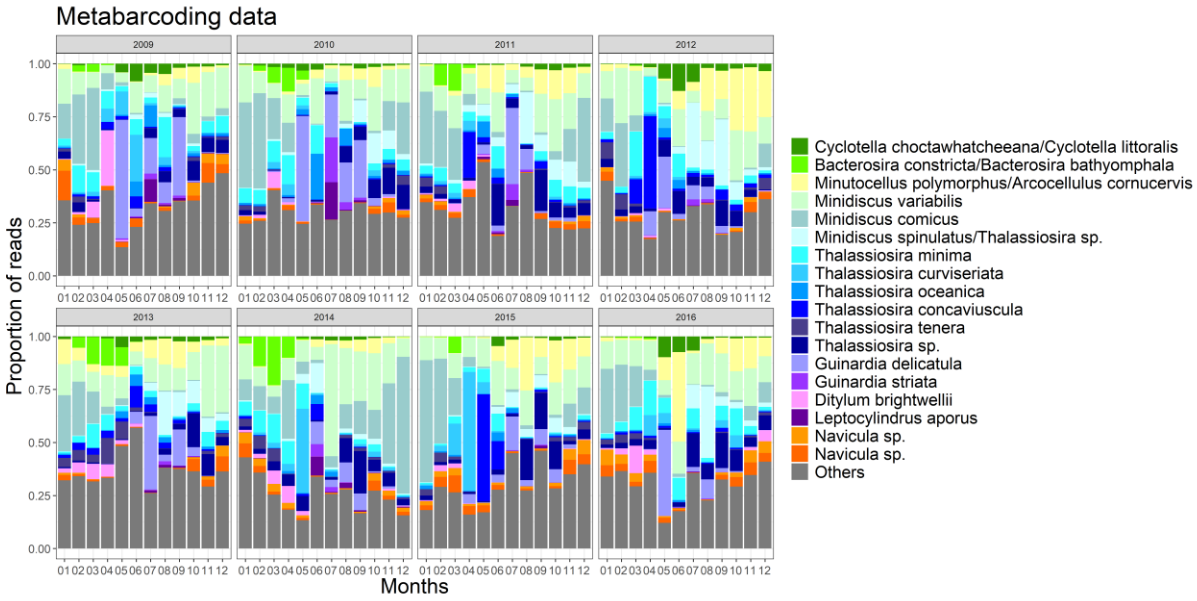
Metabarcoding data
2009
2010
2011
2012
2013
2014
2015
2016
Proportion of reads
0.00
0.25
0.50
0.75
1.00
01 02 03 04 05 06 07 08 09 10 11 12
Months
Cyclotella choctawhatcheeana/Cyclotella littoralis
Bacterosira constricta/Bacterosira bathyomphala
Minutocellus polymorphus/Arcocellulus cornucervis
Minidiscus variabilis
Minidiscus comicus
Minidiscus spinulatus/Thalassiosira sp.
Thalassiosira minima
Thalassiosira curviseriata
Thalassiosira oceanica
Thalassiosira concaviuscula
Thalassiosira tenera
Thalassiosira sp.
Guinardia delicatula
Guinardia striata
Ditylum brightwellii
Leptocylindrus aporus
Navicula sp.
Navicula sp.
Others

Table S1 : Taxonomic assignation of the 32 dominant OTUs (shaded rows) and considering only diatoms (see text for the method used) at the SOMLIT-Astan time-series station. The automatic assignation using the PR2 database was checked and refined by comparing the corresponding sequences to references using BLAST (ref). The SILVA assignations of these sequences was also checked for some dinoflagellate sequences. The origin of the best blast sequences listed (most of which had 100% similarity with our query sequence) and of the corresponding strains or isolates were carefully examined before taking the final taxonomic assignation decision. Cultured strains that were isolated from the SOMLIT-Astan Station are highlighted in bold. For some of the reference sequences, taxonomic assignation was not considered valid (assignation written in gray). Microalgae Culture Collection [1] Trefault et al. (2020), [2] Nanjappa,D. et al. (2013), [3] Laza-Martinez etal. (2012)

| PR2 taxonomic group | OTU (PR2 species rank taxonomy_OTU short name_4 first letters of OTU ID code) | Manual assignation | | | | | |
|---|---|---|---|---|---|---|---|
| | | Best blasts | Corresponding isolates or strains | % identity | NCBI Taxonomy | SILVA Taxonomy | Final assignation |
| Bacillariophyta | Guinardia_delicatula_baed | MH782132.1 | **RCC5799** | 100 | *Guinardia delicatula* | | *Guinardia delicatula* |
| Bacillariophyta | Minidiscus_a751 | MN528627.1 | **RCC4657** | 100 | *Minidiscus variabilis* | | *Minidiscus variabilis* |
| Bacillariophyta | Polar-centric-Mediophyceae_92af | LC189088; JN934677.1 | NIES-3970; RCC2270 | 100 | *Minutocellus; Arcocellulus* | | *Minutocellus/Arcocellulus* |
| Bacillariophyta | Polar-centric-Mediophyceae_dd92 | MH011734 | GF367-18S_12 | 100 | *Thalassiosira* | | *Thalassiosira sp.* |
| Bacillariophyta | Polar-centric-Mediophyceae_X_sp._38e7 | MN528601 | **RCC4660** | 100 | *Minidiscus comicus* | | *Minidiscus comicus* |
| Bacillariophyta | Thalassiosira_c57e | JN934676 | RCC2265 | 100 | *Thalassiosira minima* | | *Thalassiosira minima* |
| Bacillariophyta | Thalassiosira_cefd | MH843669.1; DQ514870 | RCC4582; CC03-15 | 100 | *Minidiscus sp.; Shionodiscus oestrupii var venrickae* | | *Minidiscus* sp.[1] |
| Bacillariophyta | Thalassiosira_379f | MN528650 | **RCC5154** | 100 | *Thalassiosira curviseriata* | | *Thalassiosira curviseriata* |
| Bacillariophyta | Thalassiosira_concaviuscula_1b87 | AJ810857 | Type strain MHtc1 | 100 | *Thalassiosira concaviuscula* | | *Thalassiosira concaviuscula* |

| | | | | | | |
|---|---|---|---|---|---|---|
| Bacillariophyta | Bacterosira_a2c6 | KT692951.1; DQ514894.1; DQ514877.1 | SMDCO1286; NB04-B6; CCMP991 | 100 | *Bacterosira constricta; Bacterosira bathyomphala; Bacterosira sp.* | *Bacterosira sp.* |
| Bacillariophyta | Cyclotella_46d6 | JQ217341.1; JQ217340.1 | KMMCC B147; KMMCC B214 | 100 | *Cyclotella choctawhatcheeana; Cyclotella litoralis* | *Cyclotella sp.* |
| Bacillariophyta | Ditylum_brightwellii_1e0f | AY485444.1; X85386.1 | CCMP358; CCAP1022/2 | 100 | *Ditylum brightwellii* | *Ditylum brightwellii* |
| Bacillariophyta | Leptocylindrus_d056 | KC814810.1; AJ535175.1 | SZN-B651; KM9950 | 100 | *Leptocylindrus aporus; Leptocylindurs danicus* | *Leptocylindrus aporus*[2] |
| Bacillariophyta | Navicula_92d5 | KT861019.1 | RCC3092 | 100 | *Navicula* | *Navicula* sp. |
| Bacillariophyta | Polar-centric-Mediophyceae_71e5 | AJ810858.1 | "type strain MHtt1" | 100 | *Thalassiosira tenera* | *Thalassiosira tenera* |
| Bacillariophyta | Radial-centric-basal-Coscinodiscophyceae_X_sp._4884 | KJ671697.1; KT861015.1 | "Strain 12"; RCC2966 | 100; 99.74 | *Guinardia striata* | *Guinardia striata* |
| Bacillariophyta | Thalassiosira_oceanica_8902 | DQ514878.1 | CCMP1001 | 100 | *Thalassiosira oceanica* | *Thalassiosira oceanica* |
| Cercozoa | Cryothecomonas_aestivalis_2a84 | AF290541.1; AF290539.1 | "Strain 2"; "Strain 1" | 99.74; 99.48 | *Cryothecomonas aestivalis* | *Cryothecomonas aestivalis* |
| Cercozoa | Cryothecomonas-lineage_7aac | KY979993.1; GQ144679.1 | NY13S_83; APCC MC-1Cryo | 99.74; 99.22 | Cryothecomonas sp. | Cryothecomonas sp. |
| Chlorophyta | Ostreococcus_lucimarinus_59f6 | MT117941.1 | BCC118000 | 100 | *Ostreococcus lucimarinus* | *Ostreococcus lucimarinus* |
| Cryptophyta | Cryptomonadales_X_39cf | MK956825.1; MK956143.1 | "strain 10"; single cell isolate | 100 | *Teleaulax amphioxeia (=Plagioselmis prolonga)* | *Teleaulax amphioxeia (=Plagioselmis prolonga)*[3] |
| Cryptophyta | Cryptomonadales_X_a5a5 | JQ966995.1; JQ966994.1 | Cr7EHU; Cr6EHU | 100 | *Teleaulax gracilis* | *Teleaulax gracilis* |
| Cryptophyta | Cryptomonadales_X_f546 | MK956814.1; HM126531.1; KY980327.1; KY980204.1 | "Strain 07"; SCAP K-1486 | 100 | *Teleaulax acuta/ Teleaulax amphioxeia* | *Teleaulax acuta*[3] |
| Cryptophyta | Cryptomonadales_XX_sp._7e7d | KY980182.1; MK956818.1 | BH46_33; RCC5152 | 99,73 | *Teleaulax amphioxeia; Geminifera cryophila* | *Teleaulax/Geminifera clade* |

| | | | | | | | |
|---|---|---|---|---|---|---|---|
| Dinoflagellata | Dinoflagellata_bcc6 | KY980212.1; KY980192.1 | BH46_94; BH46_57 | 100 | *Adenoides eludens* | *Haplozoon* sp. | *Haplozoon* sp. |
| Dinophyceae | Dinophyceae_fcf1 | KY980285.1 | BH46_144 | 100 | *Heterocapsa rotundata* | *Heterocapsa* sp. | *Heterocapsa* sp. |
| Dinophyceae | Dinophyceae_3301 | AF274267.1; DQ388464.1 | CCMP680; CCMP1542 | 100 | *Heterocapsa rotundata* | *Heterocapsa rotundata* | *Heterocapsa rotundata* |
| Dinophyceae | Dinophyceae_8fd6 | FN669511.1; FN669510.1 | GCGMS0407NS; GDMS0704YD | 99.74 | *Gyrodinium cf guturula; G. dominans* | *Gyrodinium* sp. | *Gyrodinium sp.* |
| Dinophyceae | Dinophyceae_09a4 | KY980035.1 | NY13S | 100 | *Warnowia* sp. | *Gymnodinium* clade, *Gyrodiniellum* | Gymnodinium clade |
| Dinophyceae | Dinophyceae_XXX_sp._af41 | KP790170.1; FJ947040 | 5 AR-2015; BSL-2009a | 99.21 | *Warnowia* sp. | *Warnovia* sp. | *Warnovia* sp. |
| Dinophyceae | Dinophyceae_XXX_sp._24ad | FJ024299.1 | Clone LM83 | 99.74 | *Gyrodinium helveticum* | *Gyrodinium* sp. | Gyrodinium sp. |
| Dinophyceae | Dinophyceae_XXX_sp._034f | KY980399.1 | BH65_136 | 100 | *Pentapharsodinium* sp. | *Azadinium* sp. | Azadinium sp. |
| Dinophyceae | Dinophyceae_XXX_sp._3d28 | KU640194,1; KU314867.1; EF492506.1; HM067010.1 | unnamed; unnamed; NEPCC734; MC728-D5 | 98.43 | *Ptychodiscus noctiluca; Karlodinium veneficum; Karlodinium micrum; Takayama acrotrocha* | Gymnodiniphy-cidae | *Ptychodiscus/Karenia/Bra-chidinium/Takayama clade* |
| Dinophyceae | Gyrodinium_75e1 | KY980394.1; KY980272.1; KY980220.1; AB120002.1 | BH46_129; BH46_109; BH46_107; single cell | 100 | *Heterocapsa rotundata* (3 sequences); *Gyrodinium fusiforme* | *Gyrodinium* sp. | *Gyrodinium* cf *fusiforme* |
| Eukaryota | Eukaryota_b342 | KU747081.1; KT582539.1; KM096193.1 | SZ1; NIOLM_56; LF562 | 100 | *Parengyodontium album; Halophytophtora vesicula; Engyodontium* sp. | *Parengyodon-tium* | *Parengyodontium* |
| Picozoa | Picozoa_XXX_18f8 | JX988758.1 | Unnamed culture | 100 | *Picomonas judraskeda* | | *Picomonas judraskeda* |
| Pseudofungi | MAST-1A_XX_sp._0c52 | | | | | | MAST-1A |
| Syndiniales | Dino-Group-I-Clade-1_X_sp._5e71 | | | | | | Syndiniales GroupI Clade1 |

| | | |
|---|---|---|
| Syndiniales | Dino-Group-I-Clade-1_X_sp._ac94 | Syndiniales GroupI Clade1 |
| Syndiniales | Dino-Group-II-Clade-7_X_sp._86d3 | Syndiniales GroupII Clade7 |
